# Engineering a customizable antibacterial T6SS-based platform in *Vibrio natriegens*

**DOI:** 10.1101/2021.05.04.439770

**Authors:** Biswanath Jana, Kinga Keppel, Dor Salomon

## Abstract

Bacterial pathogens are a major risk to human, animal, and plant health. To counteract the spread of antibiotic resistance, alternative antibacterial strategies are urgently needed. Here, we constructed a proof-of-concept customizable, modular, and inducible antibacterial toxin delivery platform. By engineering a type VI secretion system (T6SS) that is controlled by an externally induced on/off switch, we transformed the safe bacterium, *Vibrio natriegens*, into an effective antibacterial weapon. Furthermore, we demonstrated that the delivered effector repertoire, and thus the toxicity range of this platform, can be easily manipulated and tested. We believe that this platform can serve as a foundation for novel antibacterial bio-treatments, as well as a unique tool to study antibacterial toxins.

## INTRODUCTION

Owing to the rapid spread of multidrug-resistant bacteria and infections, as well as the lack of new effective antibiotics, we are quickly approaching a “post-antibiotic era” in which bacterial infections that are considered curable will once again be life threatening ^1,2^. Misuse and over-use of antibiotics in medicine and in animal agriculture are thought to have contributed to the emergence of antibiotic-resistant strains ^3–5^. Moreover, global environmental changes have enabled the spread of bacterial pathogens and the emergence of new pathogenic strains, as described for members of the *Vibrionaceae* family ^6^.

There is an urgent need to develop new antibacterial treatment strategies as alternatives to antibiotics to prevent and counteract the emerging threat from bacterial pathogens. Fortunately, the scientific community has accepted this challenge, and several antibacterial approaches have been developed in recent years to address this growing problem. Such approaches include, for example, phage-therapy ^7^, bacteriocins ^8,9^, customized toxins whose expression is induced upon delivery of their encoding DNA into the target pathogen ^10^, and the use of genome-editing methods, such as CRISPR-Cas, to reverse bacterial resistance to antibiotics ^11,12^. Although promising, each of these approaches has its limitations, such as reliance on specific receptors that are presented on the surface of the target cell, reliance on target cell machinery (i.e., transcription and translation) for activation, or a delivery mode against which bacteria can quickly develop resistance. Therefore, we cannot rely on a single antibacterial strategy if we wish to stay ahead in the “arms race” against bacterial pathogens.

Bacteria have been competing with each other over resources for eons ^13^; thus, they have evolved sophisticated molecular weapons to eliminate their rivals. Many Gram-negative bacteria utilize a contact-dependent, protein secretion apparatus, termed the type VI secretion system (T6SS), to deliver toxic effector proteins into neighboring cells ^14–16^. Although T6SSs were originally described as targeting eukaryotic cells ^15^, it is now clear that most of them play a role in interbacterial competition ^17^; they use brute force to deliver an arsenal of antibacterial effectors into competing, non-kin bacteria ^16^. These antibacterial effectors are encoded in operons, together with cognate immunity proteins that protect against self- or kin-intoxication by directly binding to, and occluding the effector’s active site ^18^.

T6SS’s ability to intoxicate diverse bacteria and to deliver an arsenal of toxic effectors that synergistically target conserved and essential bacterial components ^19^ has made it a lucrative, although yet largely untapped, antibacterial tool. Indeed, Ting and colleagues recently reported engineering “programmed inhibitor cells” that use surface-displayed nanobodies to specifically adhere to and intoxicate bacteria of choice in a mixed population via T6SS activity ^20^. Their proof-of-concept system relies on the constitutively active T6SS of an opportunistic pathogen, *Enterobacter cloacae*, and on its natural repertoire of antibacterial effectors.

Here, we set out to create a novel proof-of-concept bio-treatment with inducible and customizable antibacterial properties. By introducing a T6SS into *Vibrio natriegens*, we armed this safe, non-pathogenic bacterium ^21–23^ with antibacterial capabilities. We modified T6SS so that it would serve as an inducible weapon by engineering an on/off switch that allows T6SS expression only in response to an external cue. Importantly, we demonstrated that by manipulating its effector payload, it is possible to alter the antibacterial activity and the toxicity range of this platform.

## RESULTS

### Introducing an antibacterial T6SS platform into *Vibrio natriegens*

In this work, our objective was to create an antibacterial bio-treatment that utilizes the advantages of T6SS. We envisioned that such a proof-of-concept T6SS platform should: 1) be inducible and responsive to its environment; 2) be modular, thus allowing rapid customization; 3) allow the delivered effector payload to be manipulated in order to control the toxicity range; and 4) be installed in a non-pathogenic bacterium.

As a first step, we set out to determine whether we can transform a non-pathogenic bacterium lacking known antibacterial properties into an antibacterial strain by introducing an exogenous antibacterial T6SS into it. To this end, we chose *Vibrio natriegens* as the host for a T6SS-based platform; *V. natriegens* is a genetically tractable, safe marine bacterium that thrives under diverse environmental conditions ^21–23^; it does not contain an endogenous T6SS ^24^. For the T6SS-based platform, we chose *Vibrio parahaemolyticus* T6SS1 (hereafter referred to as VpT6SS1), an antibacterial T6SS that is encoded on a single gene cluster containing all the required core components for assembling a functional T6SS, as well as regulators and antibacterial effector and immunity modules ^25–27^ (Fig. 1A). In *V. parahaemolyticus*, the system is actived under warm marine-like conditions (i.e., at 30 °C and 3% NaCl) including surface sensing activation ^25^. Studies of VpT6SS1 and its homologs in several *Vibrio* strains have revealed diverse and dynamic effector repertoires that can be exploited to alter the delivered effector payload of this system ^24,26,28–31^.

**Figure 1.**
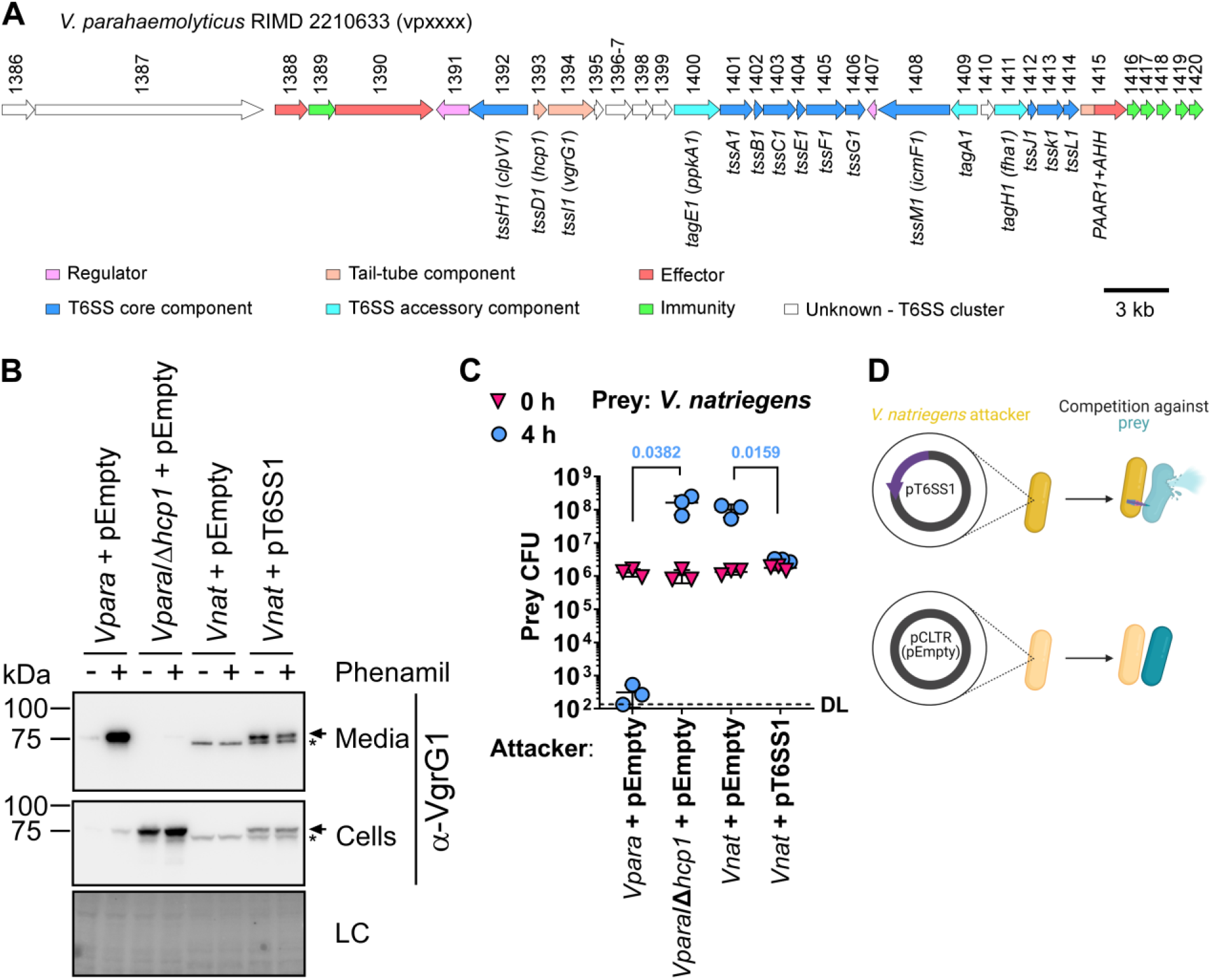
VpT6SS1 is functional in *V. natriegens*. **A)** Schematic representation of the *V. parahaemolyticus* RIMD 2210633 T6SS1 gene cluster. Genes are represented by arrows indicating the direction of transcription. Locus numbers (vpxxxx) are denoted above. Notable T6SS components are denoted below. **B)** Expression (cells) and secretion (media) of VgrG1 from *V. parahaemolyticus* RIMD 2210633 derivative POR1 (*Vpara*; T6SS1^+^) and its T6SS1^-^ mutant (*Vpara*/Δ*hcp1*), and from *V. natriegens* (*Vnat*) carrying the indicated plasmids. Samples were treated (+) or not (-) with 20 µM phenamil to activate surface sensing in media containing 3% NaCl at 30 °C for 5 h. Loading control (LC) is shown for total protein lysates. Arrows denote bands corresponding to VgrG1. Asterisks denote non-specific bands detected in *Vnat* samples. **C)** Viability counts of *V. natriegens* prey before (0 h) and after (4 h) co-incubation with the indicated attackers, as described in B, on media containing 3% NaCl at 30 °C. Data are shown as the mean ± SD. Statistical significance between samples at the 4 h timepoint by an unpaired, two-tailed Student’s *t*-test is denoted above. A significant difference was considered as *P* < 0.05. DL, assay detection limit. **D)** Illustration of interbacterial competition mediated by VpT6SS1 when introduced into *V. natriegens* (yellow) on a plasmid (pT6SS1), as shown in C. Prey cells are denoted in blue.

We cloned the VpT6SS1-encoding gene cluster from the *Vibrio parahaemolyticus* type strain RIMD 2210633 (*vp1386-vp1420*; GenBank: BA000031.2) into a low-copy plasmid (pCLTR), generating pT6SS1 (see the Materials and Methods section and Supplementary Fig. S1 for details on plasmid construction), and introduced it into *Vibrio natriegens* ATCC 14048. The plasmid-borne VpT6SS1 was functional in *V. natriegens* under warm marine-like conditions, as evident by secretion of the hallmark secreted component, VgrG1 ^15^ (Fig. 1B). As we predicted, VpT6SS1 also equipped *V. natriegens* with the ability to outcompete its parental strain under warm marine-like conditions, consequently reducing by ∼50-fold the number of viable prey bacteria remaining after 4 hours of competition (Fig. 1C-D). Although the effect on prey viability was less dramatic than that of a *V. parahaemolyticus* attacker, which carries additional VpT6SS1 effectors encoded outside of the cluster ^26^, this result indicates that a natural VpT6SS1 is functional in *V. natriegens*.

### Engineering an inducible on/off switch for VpT6SS1 in *V. natriegens*

Aiming to transform VpT6SS1 in *V. natriegens* into an inducible system that is controlled by an external cue, we sought to identify a gene that can serve as an on/off switch for VpT6SS1-mediated antibacterial activity. This required identifying a regulator whose deletion shuts off all or most of the VpT6SS1 genes; its expression should turn them back on. Three major positive regulators of VpT6SS1 have been identified previously in *V. parahaemolyticus* and were considered possible candidates: TfoY ^32,33^, which is encoded outside of the VpT6SS1 cluster, and two regulators encoded within the cluster, VP1407 and VP1391 ^25,27,33^.

Using a combination of quantitative real-time PCR (RT-PCR), interbacterial competitions, and VgrG1 secretion assays, we deciphered the role of the three regulators in the regulatory cascade of VpT6SS1 in *V. parahaemolyticus* (Supplementary Fig. S2). Importantly, we found that TfoY is an upstream regulator whose expression leads to upregulation of the *vp1409-7* operon encoding the regulator VP1407. VP1407, in turn, regulates all VpT6SS1 operons except *vp1393-9*, including its own operon. Lastly, VP1407-induced VP1391 upregulates the remaining *vp1393-9* operon to complete the VpT6SS1 activation (Fig. 2A).

**Figure 2.**
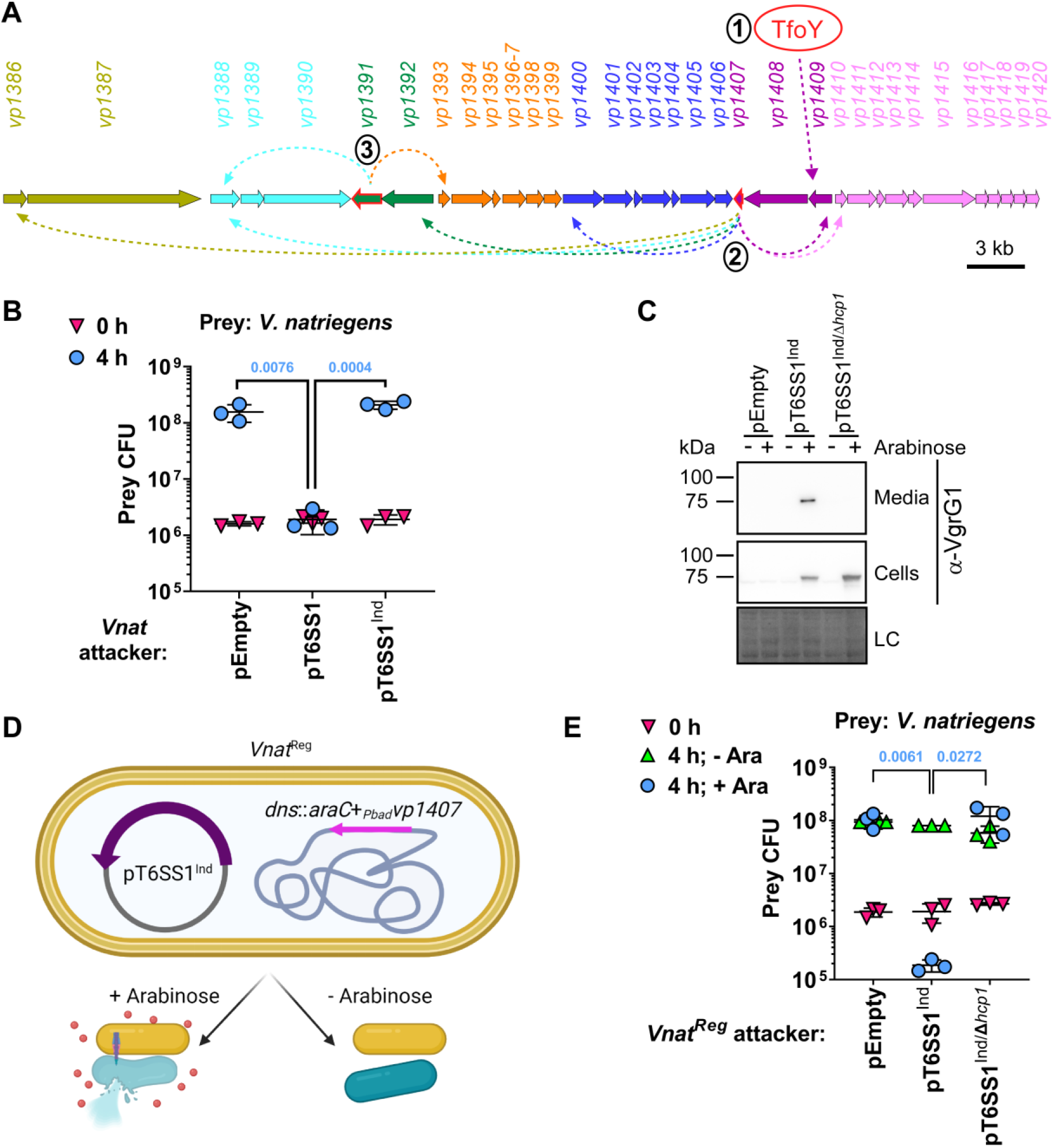
VP1407 can serve as an effective on/off switch for VpT6SS1 in *V. natriegens*. **A)** Illustration of the VpT6SS1 (*vp1386-vp1420*) activation cascade. Genes are represented by arrows indicating the direction of transcription. Genes belonging to the same operon are denoted by the same color. Locus numbers (vpxxxx) are denoted above. Positive regulators are denoted by a red frame. Dashed arrows denote transcriptional activation and are color coded according to the relevant induced operon. Numbers in black circles next to the positive regulators TfoY, VP1407, and VP1391 delineate the progression of the cascade. **B)** Viability counts of *V. natriegens* prey before (0 h) and after (4 h) co-incubation with wild type *V. natriegens* attackers harboring the indicated plasmids, on media containing 3% NaCl at 30 °C. **C)** Expression (cells) and secretion (media) of VgrG1 from *V. natriegens* containing an arabinose-inducible *vp1407* in the chromosomal *dns* locus (*Vnat*^Reg^) and harboring an empty plasmid (pEmpty) or plasmids containing VpT6SS1 that lacks *vp1407* (pT6SS1^Ind^) or both *vp1407* and *hcp1* (pT6SS1^Ind/Δ*hcp1*^; T6SS^-^). Samples were grown in media containing 3% NaCl and either supplemented (+) or not (-) with 0.1% arabinose (Ara) at 30 °C. Loading control (LC) is shown for total protein lysates. **D)** Illustration of the engineered *Vnat*^Reg^ (yellow bacteria) containing a plasmid-borne, inducible VpT6SS1 (pT6SS1^Ind^). In the presence of arabinose (red circles), VP1407 is expressed from the chromosome and the plasmid-borne T6SS is induced, resulting in T6SS-mediated intoxication of adjacent prey bacteria (blue). **E)** Viability counts of *V. natriegens* prey before (0 h) and after (4 h) co-incubation with *Vnat*^Reg^ attackers carrying the indicated plasmids on solid media plates, as described in C. For B and E, data are shown as the mean ± SD; statistical significance between samples at the 4 h timepoint by an unpaired, two-tailed Student’s *t*-test is denoted above and is color coded to match the relevant samples. A significant difference was considered as *P* < 0.05.

In *V. natriegens*, TfoY (WP_020332876.1; 89% identity to *V. parahaemolyticus* RIMD 2210633 TfoY) could not be used as a switch, since it was not required for VpT6SS1 activation (Supplementary Fig. S3). Therefore, we determined whether VP1407, whose deletion down-regulated all VpT6SS1 operons (Supplementary Fig. S2B) and whose expression induced all VpT6SS1 operons in *V. parahaemolyticus*, either directly or via VP1391 (Supplementary Fig. S2A), can serve as the on/off switch. Indeed, removing *vp1407* from the plasmid-borne VpT6SS1 (resulting in pT6SS1^Ind^) rendered the system inactive; it lost its ability to mediate interbacterial toxicity (Fig. 2B) and to express VgrG1 (Fig. 2C, no arabinose).

To function as an on/off switch, VP1407’s expression needed to be inducible. Moreover, we wanted VP1407 to be expressed *in trans* so that the platform will be modular and thus allow simple future customization. As proof-of-concept, we engineered *vp1407* into the *V. natriegens* chromosome (replacing the *dns* genomic locus ^22^) under the regulation of the arabinose-inducible *Pbad* promoter, together with the *Pbad* regulator, AraC ^34^; this resulted in a derivative strain that is hereafter termed *Vnat*^Reg^ (Fig. 2D and Supplementary Fig. S1H). As shown in Fig. 2C, VgrG1 expression and secretion were restored when *Vnat*^Reg^ carrying pT6SS1^Ind^ were grown in the presence of arabinose. Furthermore, VpT6SS1-mediated antibacterial toxicity was also restored upon arabinose addition (Fig. 2D-E). Notably, the reduction in prey viability mediated by this inducible system (∼3 orders of magnitude; Fig. 2E) was more pronounced than the reduction mediated by the natural VpT6SS1 gene cluster in *V. natriegens* (∼50-fold; Fig. 1C); these data reveal that external activation of the system can result in potent antibacterial activity. Taken together, the above results indicate that VP1407 can serve as an effective on/off switch for VpT6SS1. Importantly, we demonstrated the successful construction of a *V. natriegens* strain that can activate an engineered antibacterial platform upon sensing an external cue (Fig. 2D).

### The engineered antibacterial platform is active under a wide temperature range and against various competitors

To evaluate the usability of the engineered antibacterial platform, we determined the temperature range in which it operates. Using parental *V. natriegens* as prey, we monitored prey viability after 4 and 24 hours of competition with *Vnat*^Reg^ attackers carrying an inducible T6SS (pT6SS1^Ind^) or its inactive version (pT6SS1^Ind/Δ*hcp1*^), at temperatures ranging from 20 to 37 °C. As shown in Fig. 3A, the system was active within the tested range, although at 20 °C it required 24 h to mediate an effect similar to that seen at other temperatures within 4 h. This was probably due to the slower growth of *V. natriegens* at this temperature (Supplementary Fig. S4).

**Figure 3.**
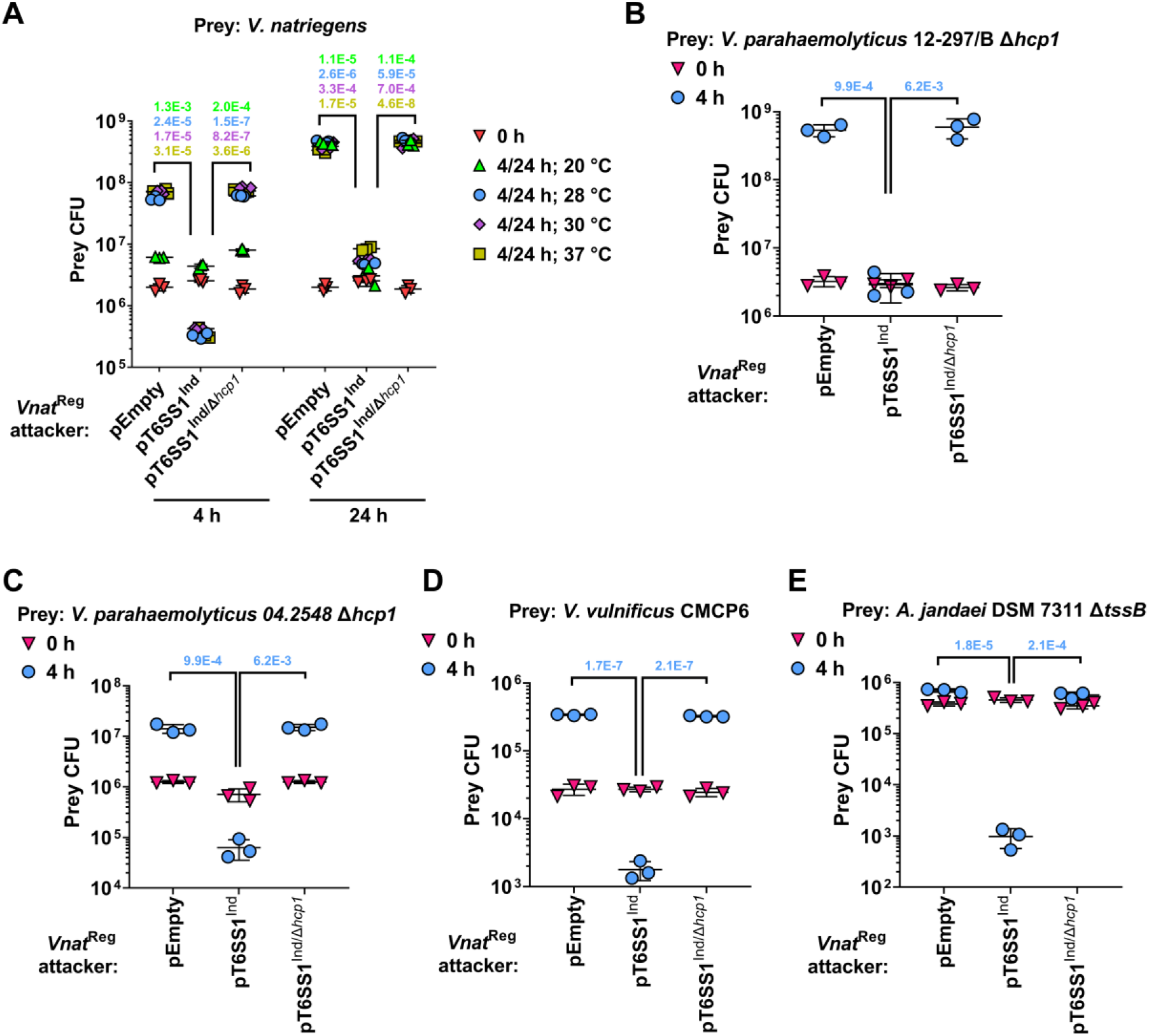
A wide-range functionality of the inducible VpT6SS1-based platform in *V. natriegens*. **A)** Viability counts of *V. natriegens* prey before (0 h) and after (4 or 24 h) co-incubation with *Vnat*^Reg^ attackers harboring the indicated plasmids, on media containing 3% NaCl and 0.1% arabinose. **B-E)** Viability counts of the indicated prey bacteria before (0 h) and after (4 h) co-incubation with *Vnat*^Reg^ attackers harboring the indicated plasmids, on media containing 3% NaCl and 0.1% arabinose at 28 °C. Data are shown as the mean ± SD. Statistical significance between samples at the 4 or 24 h timepoints by an unpaired, two-tailed Student’s *t*-test is denoted above and is color coded to match the relevant samples. A significant difference was considered as *P* < 0.05.

Because *V. natriegens* is a marine bacterium, we reasoned that as a potential bio-treatment it will be most suited to intoxicate other marine bacteria. Many marine bacteria are known or emerging pathogens of humans and aquaculture produce ^35,36^. Aquaculture produce, such as shrimp, are often farmed at temperatures around 28 °C ^37,38^, an optimum temperature at which our platform functions well (Fig. 3A). Therefore, we evaluated whether the platform can intoxicate diverse marine pathogens at this temperature. Indeed, under inducing marine-like conditions (i.e., 28 °C, 3% NaCl, and in the presence of arabinose), *Vnat*^Reg^ carrying an arabinose-inducible T6SS outcompeted pathogenic *V. parahaemolyticus* strains (the shrimp pathogen 12-297/B ^31^ and the clinical isolate 04.2548 ^39^), as well as the pathogens *V. vulnificus* and *Aeromonas jandaei* (Fig. 3B-E). Notably, to avoid masking the activity of our platform by prey-mediated counterattacks, we inactivated potentially antibacterial T6SSs in a few of the competing bacteria by deleting genes encoding the conserved T6SS components *hcp* or *tssB* ^40^, as indicated.

### Manipulating the effector repertoire of the engineered platform

One of our main aims was to control not only the activation of the antibacterial platform, but also the identity of the toxins that it deploys. The inducible T6SS platform that we have engineered still carried two endogenous antibacterial effector and immunity modules, one at each end of the cluster (*vp1388-90* and *vp1415-6*) ^26^, which mediate the intoxication of a wide range of bacteria (Fig. 3). To enable control of the effector repertoire, we first set out to remove the endogenous effectors from the platform. To this end, we constructed a version of pT6SS1^Ind^ in which *vp1388-90* have been deleted and the two histidine residues in the putative active site of the AHH toxin domain (residues 563-4), which is fused to a PAAR repeat-containing domain in VP1415, have been replaced with alanines (hereafter referred to as pT6SS1^Effectorless^). As expected, this inducible and effectorless platform was unable to mediate arabinose-inducible intoxication of a sensitive prey (Supplementary Fig. S5A). Importantly, the effectorless T6SS remained functional, as evident by the T6SS-mediated secretion of VgrG1 (Supplementary Fig. S5B) and the assembly of T6SS sheaths (Supplementary Fig. S5C).

Next, we introduced into *Vnat*^Reg^ that harbors the inducible and effectorless T6SS platform (pT6SS1^Effectorless^) various expression plasmids carrying T6SS effector and immunity modules that were placed under *Pbad* regulation. These modules were previously shown to be secreted by VpT6SS1 homologous systems (PoNe/i together with VgrG1b from *V. parahaemolyticus* 12-297/B ^28^, Tme/i1 from *V. parahaemolyticus* BB22OP ^29^, VPA1263-Vti2 from *V. parahaemolyticus* RIMD 2210633 ^26^, and Va02265-0 from *V. alginolyticus* 12G01 ^30^). As expected, the exogenous effector and immunity modules restored the platform’s ability to intoxicate parental *V. natriegens* prey. Surprisingly, however, each module differentially affected other bacteria that were used as prey (Fig. 4, left panels – “Single prey”). PoNe/i intoxicated all of the tested prey except *A. jandaei*, whereas the other three modules affected aquatic prey (i.e., *V. natriegens, V. vulnificus*, and *A. jandaei*) but not mammalian gut-residing bacteria (i.e., *E. coli* and *Salmonella enterica*) (Fig. 4). Interestingly, Tme/i1 had a major effect on vibrios, but it had only a minor effect on *A. jandaei* viability. These results show that the effector repertoire of the VpT6SS1-based platform can be manipulated. Intriguingly, these results also reveal that effectors have different toxicity ranges.

**Figure 4.**
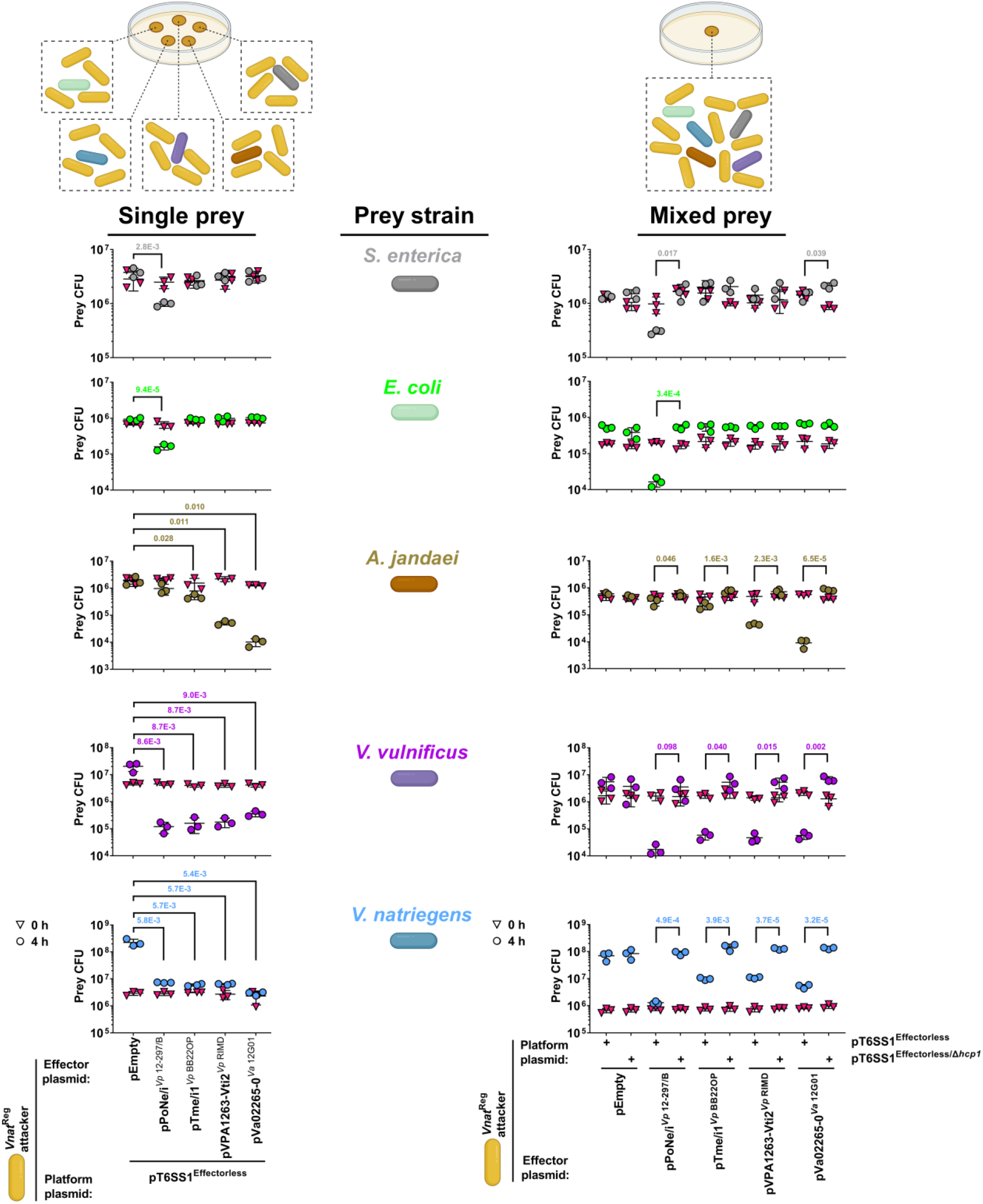
Manipulating the effector repertoire and target range of the VpT6SS1-based platform. Viability counts of *V. natriegens* (cyan), *V. vulnificus* (purple), *A. jandaei* Δ*tssB* (brown), *E. coli* (green), and *S. enterica* (gray) prey before (0 h) and after (4 h) co-incubation with *Vnat*^Reg^ attackers (yellow) harboring the indicated plasmids, on media containing 3% NaCl at 28 °C. The left panels show prey survival when competing alone (single prey) against the attacker at a 4:1 (attacker:prey) ratio; the right panels show the results of a competition experiment in which all prey were mixed together with the attacker at a 10:1:1:1:1:1 (attacker:prey) ratio. Data are shown as the mean ± SD. Statistical significance between samples at the 4 h timepoint by an unpaired, two-tailed Student’s *t*-test is denoted above and is color coded to match the relevant samples. A significant difference was considered as *P* < 0.05.

These findings prompted us to determine whether the differential toxicity of the tested effectors can enable selective targeting of specific bacteria within a mixed population. Indeed, the same phenomenon was observed when the five above-mentioned prey strains were all mixed together and competed against our platform (Fig. 4, right panels – “Mixed prey”), indicating that natural, exogenously expressed effectors can be used to target specific bacteria within a diverse prey community via T6SS activity.

### Deploying multiple effectors by the engineered platform

Lastly, we set out to demonstrate that the engineered platform can deliver multiple exogenous effectors, and to determine the advantage of deploying multiple effectors to widen the platform’s target range. To this end, we engineered a plasmid expressing a combination of two effectors: VPA1263 ^26^ from *V. parahaemolyticus* RIMD 2210633, and Va02265 ^30^ from *V. alginolyticus* 12G01, together with their cognate immunity genes to prevent self-intoxication. As expected, single effectors could mediate the intoxication of a prey that was not the strain from which they were derived. Nevertheless, their combination mediated T6SS-dependent intoxication of both prey strains (Fig. 5A-C). Notably, antibacterial T6SSs in the tested prey strains were inactivated (by deleting the conserved component *hcp*) to prevent counterattacks that could mask the effect of the engineered platform. Surprisingly, combining the two effector and immunity modules did not provide a significant advantage over a single module, when deployed against a prey that is sensitive to both effectors (i.e., *V. natriegens*) (Fig. 5D). These results demonstrate the ability of the engineered platform to deploy multiple effectors, as well as the applicability of using multiple effectors to widen its toxicity range.

**Figure 5.**
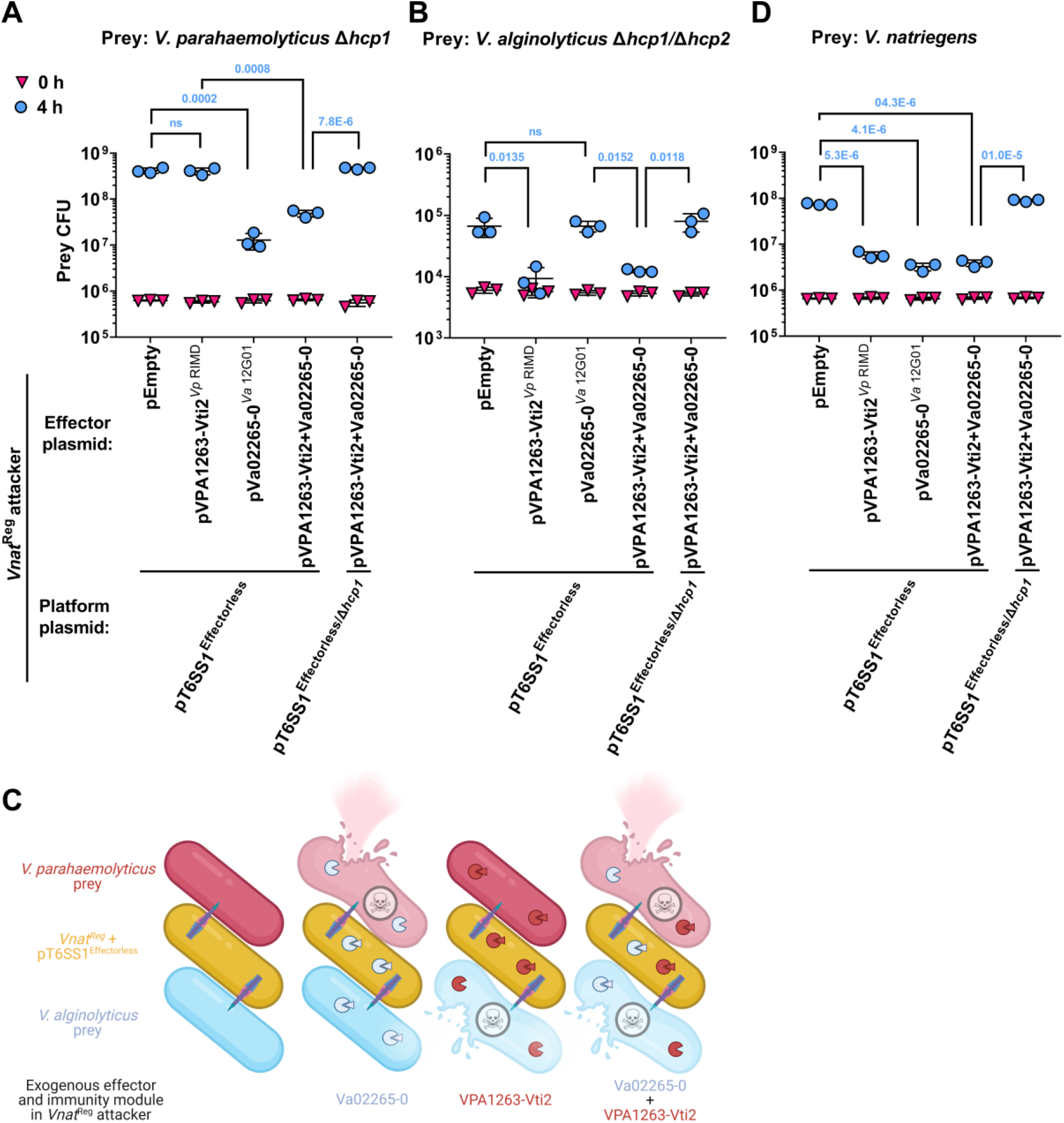
Effector and immunity combinations alter the VpT6SS1-based platform’s toxicity range. **A-B and D)** Viability counts of *V. parahaemolyticus* RIMD 2210633 Δ*hcp1* (A), *V. alginolyticus* 12G01 Δ*hcp1*/Δ*hcp2* (B), and *V. natriegens* (D) prey bacteria before (0 h) and after (4 h) co-incubation with *Vnat*^Reg^ attackers harboring the indicated plasmids, on media containing 3% NaCl and 0.1% arabinose at 28 °C. Data are shown as the mean ± SD. Statistical significance between samples at the 4 h timepoint by an unpaired, two-tailed Student’s *t*-test is denoted above. A significant difference was considered as *P* < 0.05. **C)** Illustration summarizing the competitions shown in A-B. Effector and immunity modules (denoted as a circle and a triangle, respectively) expressed by *Vnat*^Reg^ carrying pT6SS^Effectorless^ (yellow) are colored to match the bacterium from which they were derived.

## DISCUSSION

This work presents a proof-of-concept engineered, modular bacterial platform for the controlled delivery of antibacterial effectors via T6SS. We propose that this platform can serve as a foundation for custom-designed, synthetic bio-treatments against bacterial pathogens. One of the platform’s main advantages lies in its customizability; it can be engineered to respond to an external cue by modifying the regulation of an on/off switch, and its effector arsenal can also be manipulated. Moreover, an important attribute of this platform is its modularity. The effectorless T6SS-encoding gene cluster, which is inactive by itself (it lacks *vp1407*), is separated from the on/off switch and from the effector and immunity modules, both of which are provided *in trans*. This modularity simplifies future steps aimed at customizing the regulation of the platform and its effector repertoire, and thus its potential target range. Furthermore, this modular design takes into account the need to prevent future acquisition of this engineered system by other bacteria upon its deployment as a bio-treatment; the physical separation of its components should hinder acquisition of a complete and functional system by competitors via horizontal gene transfer.

We envision that this platform can be used as a specific pathogen-targeted bio-treatment upon further customization, taking under consideration the following themes: 1) the on/off switch should be induced upon sensing the pathogen to be targeted or the relevant niche, thus preventing purposeless activation of an energy-consuming apparatus. This can be achieved, for example, by appropriating the pathogen-of-target’s quorum sensing regulation machinery to drive the expression of the platform’s on/off switch; 2) the platform should be able to intoxicate only desired pathogens while remaining benign to other bacteria in order to avoid unwanted dysbiosis. This can be achieved by equipping it with effectors that exhibit target-specific toxicity, either natural or synthetic. Alternatively, specificity can be achieved by integrating this platform with adhesion-mediated targeting mechanisms. Ting et al. recently described such a method, in which a bacterium with antibacterial capabilities is selectively targeted to a pathogen of interest within a mixed community by surface-displayed nanobodies that mediate specific cell-cell adhesion ^20^; 3) the effector and immunity modules that will be introduced into the platform should be directly regulated by the T6SS on/off switch. This can be achieved by having them controlled by a promoter from a VpT6SS1 operon that is directly upregulated by VP1407; 4) a deployed bio-treatment should be safe. Here, the platform is hosted by *V. natriegens*, which is considered to be a safe, non-pathogenic bacterium ^23^. Nevertheless, prior to its deployment, even this bacterium should be modified to limit its ability to acquire external genetic information that may include virulence factors, e.g., by hindering its natural competency machinery; 5) the deployed platform should be stable. Notably, in its current proof-of-concept state, parts of the platform are plasmid-borne. In future deployable versions, all of the modules should be introduced into the bacterial chromosome to ensure their long-term stability.

The current host for the platform, *V. natriegens*, can survive and grow under diverse conditions ^22^. Since it is a natural inhabitant of marine environments ^41^, we predict that it can serve as a bio-treatment in marine aquaculture settings. Towards this goal, we demonstrated that the platform is functional under conditions suitable for aquaculture farming, and that it is active against diverse marine pathogens. We predict that this or similar platforms can be integrated into diverse bacterial hosts that thrive in different environments, thus creating T6SS-based synthetic bio-treatments that can be used in an assortment of environmental and clinical setups.

In addition to its possible applicability as a bio-treatment, the effectorless platform also serves as a valuable research tool that can provide answers to basic questions in the T6SS field. For example, since single effectors can be introduced into the effectorless platform, the antibacterial effect of the attacking strain will be mediated solely by one effector. Therefore, the effector can be studied under the natural conditions in which it is delivered into diverse prey cells by T6SS, instead of being exogenously over-expressed inside a surrogate cell. Furthermore, the delivery of single effectors permits an analysis of the prey factors responsible for sensitivity or resistance. Here, we found that effectors can intoxicate certain bacterial prey, whereas they have no effect on other bacteria. Although this phenomenon could be useful to provide our platform with a narrow target range against specific bacterial pathogens, it also underscores the use of diverse effector arsenals by natural T6SSs to ensure intoxication of various competitors. However, it remains to be determined whether the observed lack of toxicity against certain bacteria is a property of an effector’s activity, the ability of the T6SS to deliver the effector into certain prey, the environmental conditions under which the competition took place, or a property of the prey that renders it resistant to the activity of certain effectors. Notably, natural resistance of bacteria against T6SS effector intoxication, which is not mediated by cognate immunity proteins, has been recently reported in other studies ^42–45^. Therefore, our platform serves as a unique tool that allows T6SS-mediated secretion of single effectors under a wide range of environmental conditions, enabling one to study effector toxicity ranges and revealing natural resistance mechanisms and strategies.

This platform can also be used to study effector synergy, a concept that was recently proposed by LaCourse et al. ^19^. It allows deployment of multiple effectors of choice, and can be competed against different prey strains under diverse environmental conditions. The data that can be mined from such analyses, together with the above-mentioned effector target specificity, can be used when constructing a pathogen-oriented effector repertoire for a deployable bio-treatment.

In conclusion, we describe here the engineering of a bacterial platform that can be used to develop antibacterial bio-treatments. Upon future adaptation of its activation cues and optimization of its effector repertoire to allow specific recognition and targeting of a bacterial pathogen-of-interest, respectively, the efficiency of this platform and its ability to protect against bacterial infections will be tested *in vivo*.

## MATERIALS AND METHODS

### Strains and Media

For a complete list of strains used in this study, see Supplementary Table S1. *Vibrio parahaemolyticus* and *Vibrio natriegens* were routinely grown in MLB media (Lysogeny broth [LB] containing 3% wt/vol NaCl) or on marine minimal media (MMM) agar plates (1.5% wt/vol agar, 2% wt/vol NaCl, 0.4% wt/vol galactose, 5 mM MgSO_4_, 7 mM K_2_SO_4_, 77 mM K_2_HPO_4_, 35 mM KH_2_PO_4_, and 2 mM NH_4_Cl) at 30 °C. *Vibrio alginolyticus* were grown in MLB media and on MLB agar (1.5% wt/vol) plates at 30 °C. *Vibrio vulnificus, Aeromonas jandaei*, and *Salmonella enterica* were grown in LB media or on LB agar (1.5% wt/vol) plates. *V. vulnificus* and *A. jandaei* were grown at 30 °C, whereas *S. enterica* was grown at 37 °C. When *V. parahaemolyticus, V. alginolyticus, A. Jandaei*, or *V. natriegens* harbored a plasmid, chloramphenicol (10 μg/mL), kanamycin (250 μg/mL), or gentamycin (50 μg/mL) was added to the media to maintain the plasmid. *E. coli* were grown in 2xYT broth (1.6% wt/vol tryptone, 1% wt/vol yeast extract, and 0.5% wt/vol NaCl) at 37 °C. When *E. coli* harbored a plasmid, chloramphenicol (10 μg/mL), kanamycin (30 μg/mL), erythromycin (250 μg/mL), or gentamycin (50 μg/mL) was added to the media to maintain the plasmid. When the expression of genes from an arabinose-inducible promoter was required, L-arabinose was added to the media at 0.1% wt/vol, as specified.

*Saccharomyces cerevisiae* MaV203 yeast were grown on yeast synthetic defined (SD) agar plates (0.67% wt/vol yeast nitrogen base without amino acids, 0.14% wt/vol yeast synthetic drop-out medium supplement, 2% wt/vol glucose, 0.01% wt/vol leucine, 0.002% wt/vol uracil, 0.002% wt/vol histidine, 0.002% wt/vol tryptophan, and 2% wt/vol agar) at 30 °C.

### Plasmid construction

For a complete list of plasmids used in this study, see Supplementary Table S2. Primers used for amplification are listed in Supplementary Table S3.

To construct pCLTR, a conjugatable, low-copy *E. coli*-yeast-*Vibrio* shuttle vector, a fragment consisting of the *p15A ori* and *Cm*^*R*^ genes from pBAD33, and *oriT* from pUC18T-mini-Tn7T-Tp-dsRedExpress were PCR amplified and introduced into a linearized pYES1L plasmid (Novagen) using the Gibson assembly method ^46^ (Supplementary Fig. S1A).

For arabinose-inducible expression, the coding sequences (CDS) of *vp1391* and *vp1407* were amplified from *V. parahaemolyticus* POR1 genomic DNA and inserted into the multiple cloning site (MCS) of pBAD/*Myc*-His (Invitrogen) vector containing a kanamycin-resistance cassette ^25^ (in-frame or out-of-frame with the C-terminal *Myc*-His tag, respectively), using the Gibson assembly method, to generate pVP1391 and pVP1407, respectively. The construction of pTfoY was described previously ^33^.

For arabinose-inducible expression of TssB1-sfGFP or of effector and immunity modules in *V. natriegens*, these cassettes were first introduced into the above-mentioned pBAD/*Myc*-His. Effector and immunity modules were amplified from their respective encoding bacterium. For TssB1-sfGFP, *tssB1* (*vp1402*) and *sfgfp* were amplified from *V. parahaemolyticus* RIMD 2210633 genomic DNA and sfGFP-N1 (Addgene), respectively. The amplified cassettes were inserted into the pBAD/*Myc*-His MCS using the Gibson assembly method so that the 3’ end is in-frame with the C-terminal *Myc*-His tag in the plasmid, except for Va02265-0 from *V. alginolyticus* 12G01. TssB1 and sfGFP were separated by a six-amino acid-long linker (AAAGGG). Next, the cassettes were amplified from pBAD/*Myc*-His (the region spanning the P*bad* promoter to the *rrnT1* terminator) and inserted into pVSV209 ^47^, replacing its *rfp, Cm*^*R*^, and *gfp* genes, using the Gibson assembly method (Supplementary Fig. S1B). The resulting plasmids were pPoNe/i^*Vp* 12-27/B^, pTme/i1^*Vp* BB22OP^, pVPA1263-Vti2^*Vp* RIMD^, pVa02265-0^*Va* 12G01^, and pTssB1-sfGFP (collectively termed pEI-x in Supplementary Fig. S1B). To construct a plasmid with two effector and immunity modules (pVPA1263-Vti2+Va02265-0), the above-mentioned cassette containing Va02265-0 was amplified from pBAD/*Myc*-His (the region spanning the P*bad* promoter to the *rrnT1* terminator) and inserted into pVPA1263-Vti2^*Vp* RIMD^ at the 3’ and of the VPA1263-Vti2 cassette. As a control plasmid for these pVSV209-derived vectors, we used pBJ209-araC, which is a pVSV209 in which the *rfp-gfp* region was replaced with the region spanning the *araC* CDS to the *rnnT1* terminator from pBAD/*Myc*-His, using the Gibson assembly method.

pT6SS1, and its derivatives pT6SS1^Ind^, pT6SS1^Ind/Δ*hcp1*^, pT6SS1^Effectorless^, and pT6SS1^Effectorless/Δ*hcp1*^ were constructed using the GeneArt High-Order Genetic Assembly System kit (Invitrogen), following the manufacturer’s instructions with minor modifications. Briefly, VpT6SS1 (*vp1386*-*vp1420*) was divided into 6.5-10 kb fragments with overlapping ends (40-150 bp) to allow their union via homologous recombination in yeast. The overlapping fragments were PCR amplified individually from *V. parahaemolyticus* genomic DNA. Next, 200 ng of each PCR fragment were added to 200 ng of a linearized pCLTR backbone, after purification from an agarose gel using the Gel/PCR DNA Fragments Extraction kit (Geneaid), and the DNA mix was precipitated using the sodium acetate and ethanol method and washed twice with 70% ethanol. After having been air dried, the precipitated DNA was resuspended in 10 µL of Milli-Q-treated ultra-pure water and mixed with competent MaV203 yeast cells supplied with the kit. Transformed yeast were plated on SD agar plates lacking tryptophan, and then incubated for 2-3 days at 30 °C. Colony PCR was performed on selected colonies to identify positive yeast colonies containing the desired construct. A positive yeast colony was scraped off a plate and lysed; then the lysate was used for electroporation into *E. coli* DH5α (λ pir). Lastly, the plasmid was transferred into *V. natriegens* via conjugation. To construct pT6SS1, *V. parahaemolyticus* POR1 genomic DNA was used as a template and the pCLTR backbone was linearized by restriction digestion with the I-SceI restriction enzyme (Supplementary Fig. S1C). For pT6SS1^Ind^, *V. parahaemolyticus* POR1 Δ*vp1407* ^25^ genomic DNA was used as a template (Supplementary Fig. S1D); for pT6SS1^Ind/Δ*hcp1*^, both *V. parahaemolyticus* POR1 Δ*vp1407* and *V. parahaemolyticus* POR1 Δ*hcp1* ^25^ were used as templates (Supplementary Fig. S1E); for pT6SS1^Effectorless^, both *V. parahaemolyticus* POR1 Δ*vp1407* and *V. parahaemolyticus* POR1 *vp1415*^AAA^ (see the description below) were used as templates (Supplementary Fig. S1F); for pT6SS1^Effectorless/Δ*hcp1*^, *V. parahaemolyticus* POR1 Δ*vp1407, V. parahaemolyticus* POR1 Δ*hcp1*, and *V. parahaemolyticus* POR1 *vp1415*^AAA^ were used as templates (Supplementary Fig. S1G). For the four pT6SS1 derivatives, the linearized pCLTR backbone was PCR amplified from pT6SS1 to include the ends of VpT6SS1 so that longer regions of identity to the cluster fragments would be available for recombination.

### Construction of deletion and mutant strains

To construct a *V. parahaemolyticus* POR1 derivative in which the putative active site of the VP1415 toxin is mutated (*vp1415*^AAA^), we first PCR amplified a region spanning 1.1 kb upstream and 1.1 kb downstream of the AHH motif (amino acids 562-4 in NP_797794.1) and inserted it into the MCS of the suicide vector pDM4 ^48^ (generating pDM4:*vp1415*), using the Gibson assembly method. We then used site-directed mutagenesis to replace the codons encoding histidines 563-4 with alanines, to generate pDM4:*vp1415*^AAA^. pDM4:*vp1415*^AAA^ was introduced into *V. parahaemolyticus* POR1 via tri-parental mating, and trans-conjugants were selected on MMM agar plates supplemented with chloramphenicol (10 µg/mL). The resulting trans-conjugants were grown on MMM agar plates containing sucrose (20% wt/vol) for counterselection and loss of the *sacB*-containing pDM4. The resulting *V. parahaemolyticus* POR1 *vp1415*^AAA^ strain was confirmed by amplifying and sequencing the relevant genomic region.

The construction of in-frame deletions of *vp1391, vp1407, tfoY* (*vp1028*), and *hcp1* (*vp1393*) in *V. parahaemolyticus* POR1, *hcp1* (*b5c30_rs15290*) in *V. parahaemolyticus* 12-297/B, and *hcp1* (*v12g01_01540*) and *hcp2* (*v12g01_07583*) in *V. alginolyticus* 12G01 were described previously ^25,28,30,33^. The triple mutant *V. parahaemolyticus* POR1 Δ3xRegulators (Δ*vp1391*/Δ*vp1407*/Δ*tfoY*) strain was constructed using the same pDM4 plasmids used previously to generate the single deletion mutants (pDM4:*vp1391*, pDM4:*vp1407*, and pDM4:*tfoY*). Deletions was performed sequentially, as previously described ^25^.

For in-frame deletion of *V. natriegens tfoY* (*m272_rs24650*), *V. parahaemolyticus* 04.2548 *hcp1* (*ba740_rs16850*), and *A. jandaei* DSM 7311 *tssB* (*bn1126_rs13720*), their respective 1 kb upstream and 1 kb downstream regions were cloned into the pDM4 MCS using restriction enzyme digestion and ligation. Plasmids were introduced into the respective strains and deletions were performed as previously described ^25^.

To construct *Vnat*^Reg^, the 1 kb upstream and the 1 kb downstream of *V. natriegens dns* gene (*pn96_00865*) were first cloned into pDM4 using restriction digestion and ligation, to generate pDM4:*dns*^*Vnat*^, into which sequences that we used to replace *dns* could be introduced. Next, the regions spanning the *araC* CDS to the *rrnT1* terminator were PCR amplified from pBAD/*Myc*-His plasmids into which *vp1407* or *vp1409-7* were introduced using the Gibson assembly method to generate pVP1407 and pVP1409-7, respectively. The amplified regions were inserted into pDM4:*dns*^*Vnat*^ between the *dns* 1 kb upstream and 1 kb downstream regions, using restriction digestion and ligation, to generate pDM4:*dns*^*Vnat*^_*vp1407* and pDM4:*dns*^*Vnat*^_*vp1409-7*, respectively. Next, pDM4:*dns*^*Vnat*^_*vp1409-7* was introduced into *V. natriegens* via tri-parental mating, and trans-conjugates in which the *dns* CDS had been replaced by the *araC*+*vp1409-7* cassette were obtained as described above (generating *Vnat*^*dns::araC+vp1409-7*^). Finally, we introduced pDM4:*dns*^*Vnat*^_*vp1407* into *Vnat*^*dns::araC+vp1409-7*^ and selected for trans-conjugates in which the *araC*+*vp1407* cassette was present between the *dns* 1 kb upstream and 1 kb downstream regions in the chromosome (generating *Vnat*^Reg^) (Supplementary Fig. S1H).

### Bacterial competition assays

Bacterial competitions were performed as described previously ^25^. Briefly, attacker and prey strains were grown overnight in liquid media (MLB at 30 °C for *V. parahaemolyiticus, V. natriegens*, and *V. alginolyticus*; 2xYT at 37 °C for *E. coli*; LB at 30 °C for *V. vulnificus* and *A. jandaei*, and at 37 °C for *S. enterica*), supplemented with antibiotics when maintenance of plasmids was required. Cultures were normalized to OD_600_ = 0.5 and were mixed at a 4:1 (attacker:prey) ratio when only a single prey strain was used. For bacterial competitions in which 5 different prey strains were mixed together, the attacker and prey were mixed at a 10:1:1:1:1:1 (Attacker:*E. coli*:*V. vulnificus*:*A. jandaei*:*S. enterica*:*V. natriegens*) ratio. Triplicates of each competition mixture were spotted on LB, MLB, or MLB supplemented with 0.1% (w/v) L-arabinose (when expression from *Pbad* promoter was required) agar plates and incubated for 4 h or 24 h at 23, 28, 30, or 37 °C, as indicated. Prey viability was determined as colony forming unit (CFU) counts at the indicated timepoints. When necessary, prey strains contained plasmids to provide a selection marker (for *V. natriegens*: pBAD/Myc-His, pBAD18, or pCLTR; for *V. parahaemolyticus, V. alginolyticus*, and *A. jandaei*: pBAD18; for *E. coli*: pBAD18 or pTnp1222). Assays were repeated two to three times with similar results; the results from a representative experiment are shown.

### VgrG1 expression and secretion assay

Expression and secretion of VgrG1, a hallmark secreted protein of VpT6SS1, were determined as described previously ^31^, with minor modifications. The indicated *V. parahaemolyticus* and *V. natriegens* strains were grown overnight in MLB media supplemented with appropriate antibiotics to maintain the plasmids. Cultures were normalized to OD_600_ = 0.18 in the same media, and L-arabinose (0.1% w/v) was added to induce expression from *Pbad* promoters, where indicated. 20 µM of Phenamil (an inhibitor of the polar flagella used to mimic surface sensing activation ^25^) was added to induce T6SS1, where indicated. Normalized cultures were grown for 5 h (for *V. parahaemolyticus*) or 3.5 h (for *V. natriegens*, unless otherwise stated) at 30 °C or 28 °C, as indicated, with constant shaking. To determine the expression of VgrG1 (cells), 1.0 OD_600_ units were pelleted and cells were resuspended in 2x Tris-glycine SDS sample buffer (Novex, Life Sciences). For secreted fractions (media), 10 OD_600_ units were pelleted and the supernatants were filtered (0.22 µm). Proteins were precipitated from the supernatants using sodium deoxycholate and trichloro acidic acid ^49^. Protein precipitates were washed twice with cold acetone and air-dried. Finally, the precipitates were resuspended in 20 µL of 150 mM Tris-Cl (pH=8.0), followed by the addition of 20 µL of 2x Tris-glycine SDS sample buffer (Novex, Life Sciences). Expression and secretion samples were incubated at 95 °C for 10 minutes and then resolved on TGX Stain-free gel (Bio-Rad). Next, proteins were transferred onto nitrocellulose membranes using Trans-Blot Turbo Transfer (Bio-Rad). Membranes were immunoblotted with custom-made anti-VgrG1 antibodies ^31^ at 1:1000 dilution. The loading of total protein lysates was visualized by analyzing trihalo compounds’ fluorescence of the immunoblot membrane. Protein signals were visualized using an enhanced chemiluminescence (ECL) substrate. The experiments were repeated two to three times with similar results. Results from a representative experiment are shown.

### Quantitative RT-PCR

*V. parahaemolyticus* isolates were grown overnight in MLB. The media were supplemented with antibiotics when it was appropriate to maintain plasmids. Overnight cultures were normalized to OD_600_ = 0.18 in 5 mL MLB and then grown at 30 °C for 2 h. Phenamil (20 µM) was added to induce the expression of the VpT6SS1 genes ^25^. When required, media were supplemented with 0.1% (w/v) L-arabinose to express plasmid-encoded genes regulated by P*bad* promoters. Cells equivalent to 1.0 OD_600_ units were harvested and washed with RNAprotect Bacteria Reagent (Qiagen). Next, RNA was isolated from the pelleted bacteria using the Bacterial RNA kit (Biomiga), following the manufacturer’s instructions. Complementary DNA (cDNA) was synthesized from isolated RNA (1 µg) using the SuperScript III First-Strand Synthesis System for RT-PCR kit (Invitrogen), following the manufacturer’s instructions. Random hexamer primer mix (Invitrogen) was used for cDNA synthesis. For RT-PCR, 2 ng template cDNA were mixed (in triplicate) with forward and reverse primers (300 nM each) and with 2x Fast SYBR Green Master Mix (Applied Biosystems). The RT-PCR and analysis were carried out using a QuantStudio 12K Flex instrument and software (Applied Biosystems). 16s rRNA was used as the endogenous control and the differential gene expression, reported as the fold change, were analyzed by the 2^-ΔΔCt^ method. Primer sets were designed using the Primer3web server ^50^ to amplify a representative gene from each VpT6SS1 cluster operon (i.e., *vp1386, vp1388, vp1392, vp1393, vp1406, vp1409*, and *vp1414*, as detailed in Supplementary Table S3).

### Bacterial growth assays

Overnight-grown cultures of *V. natriegens* were normalized to an OD_600_ of 0.01 in MLB media and transferred to 96-well plates (200 µL per well; n = 4). Cultures were grown at the indicated temperatures in a BioTek SYNERGY H1 microplate reader with constant shaking at 205 cpm. OD_600_ readings were acquired every 10 min.

### T6SS sheath assembly

pTssB1-sfGFP, a plasmid for the arabinose-inducible expression of the VpT6SS1 sheath protein TssB1 (VP1402) fused to sfGFP, was introduced via conjugation into *Vnat*^Reg^ strains carrying pT6SS1^effectorless^ or pT6SS1^effectorless/Δ*hcp1*^. Bacteria were grown overnight in MLB media supplemented with appropriate antibiotics to maintain the plasmids, and cultures were then normalized to OD_600_ = 0.18 in MLB media supplemented with antibiotics and 0.1% (w/v) L-arabinose. Normalized cultures were grown for 3 h at 28 °C, and 100 µL of each culture were harvested and washed with M9 media twice. Next, cells were resuspended in 100 µL of M9 media, and 1 µL of bacterial suspensions were spotted onto MLB-agarose (1.5% w/v) pads supplemented with 0.1% (w/v) L-arabinose. Pads were allowed to dry for 5 minutes and then placed face-down in 35mm glass-bottom Cellview cell culture dishes. Bacteria were imaged in a Nikon Eclipse Ti2-E inverted motorized microscope equipped with a CFI PLAN apochromat DM 100X oil lambda PH-3 (NA, 1.45) objective lens, a Lumencor SOLA SE II 395 light source, and ET-EGFP (#49002, used to visualize the GFP signal) filter sets. Images were acquired using a DS-QI2 Mono cooled digital microscope camera (16 MP), and were postprocessed using Fiji ImageJ suite ^51^.

### Statistical analysis

Data were analyzed with GraphPad Prism 9 and Microsoft Excel, using unpaired, two-tailed Student’s *t*-test, unless otherwise indicated. Differences of *P <* 0.05 were considered significant.

## ACKNOWLEDGMENTS

This project received funding from the European Research Council (ERC) under the European Union’s Horizon 2020 research and innovation program (Grant agreement No. 714224), and the Israel Science Foundation (ISF; grant no. 920/17) to DS. We thank members of the Salomon lab for technical assistance and helpful discussions. We also wish to thank Udi Qimron and Eran Bosis for their critical reading of the manuscript, as well as Karla Satchell, Swapan Banerjee, Udi Qimron, and Ohad Gal-Mor for providing bacterial strains, and Eric V. Stabb for providing bacterial strains and plasmids. Panels in Figures 1, 2, 4, and 5, and in Supplementary Figure S1 were created using BioRender.com.

## REFERENCES

1. World Health Organization. Antibiotic resistance. https://www.who.int/news-room/fact-sheets/detail/antibiotic-resistance (2018).

2. WHO & World Health Organization. Antimicrobial resistance. Global Report on Surveillance. World Health Organization 232 (2014). doi:10.1007/s13312-014-0374-3.

3. Martin, M. J., Thottathil, S. E. & Newman, T. B. Antibiotics overuse in animal agriculture: A call to action for health care providers. American Journal of Public Health vol. 105 2409–2410 (2015).

4. Manyi-Loh, C., Mamphweli, S., Meyer, E. & Okoh, A. Antibiotic use in agriculture and its consequential resistance in environmental sources: Potential public health implications. Molecules vol. 23 (2018).

5. Shallcross, L. J. Editorials: Antibiotic overuse: A key driver of antimicrobial resistance. British Journal of General Practice vol. 64 604–605 (2014).

6. Vezzulli, L. et al. Climate influence on Vibrio and associated human diseases during the past half-century in the coastal North Atlantic. Proc. Natl. Acad. Sci. 113, E5062–E5071 (2016).

7. Furfaro, L. L., Payne, M. S. & Chang, B. J. Bacteriophage therapy: clinical trials and regulatory hurdles. Front. Cell. Infect. Microbiol. 8, 376 (2018).

8. Kumariya, R. et al. Bacteriocins: Classification, synthesis, mechanism of action and resistance development in food spoilage causing bacteria. Microb. Pathog. 128, 171–177 (2019).

9. Yang, S. C., Lin, C. H., Sung, C. T. & Fang, J. Y. Antibacterial activities of bacteriocins: application in foods and pharmaceuticals. Front. Microbiol. 5, 241 (2014).

10. López-Igual, R., Bernal-Bayard, J., Rodríguez-Patón, A., Ghigo, J.-M. & Mazel, D. Engineered toxin–intein antimicrobials can selectively target and kill antibiotic-resistant bacteria in mixed populations. Nat. Biotechnol. 37, 755–760 (2019).

11. Goren, M., Yosef, I. & Qimron, U. Sensitizing pathogens to antibiotics using the CRISPR-Cas system. Drug Resist. Updat. 30, 1–6 (2017).

12. Pursey, E., Sünderhauf, D., Gaze, W. H., Westra, E. R. & van Houte, S. CRISPR-Cas antimicrobials: Challenges and future prospects. PLOS Pathog. 14, e1006990 (2018).

13. Hibbing, M. E., Fuqua, C., Parsek, M. R. & Peterson, S. B. Bacterial competition: surviving and thriving in the microbial jungle. Nat. Rev. Microbiol. 8, 15–25 (2010).

14. Mougous, J. D. et al. A virulence locus of Pseudomonas aeruginosa encodes a protein secretion apparatus. Science. 312, 1526–1530 (2006).

15. Pukatzki, S. et al. Identification of a conserved bacterial protein secretion system in Vibrio cholerae using the Dictyostelium host model system. Proc. Natl. Acad. Sci. 103, 1528–1533 (2006).

16. Jana, B. & Salomon, D. Type VI secretion system: a modular toolkit for bacterial dominance. Future Microbiol. 14, fmb-2019-0194 (2019).

17. Hood, R. D., Peterson, S. B. & Mougous, J. D. From striking out to striking gold: discovering that Type VI secretion targets bacteria. Cell Host Microbe 21, 286–289 (2017).

18. Alcoforado Diniz, J., Liu, Y. C. & Coulthurst, S. J. Molecular weaponry: Diverse effectors delivered by the Type VI secretion system. Cell. Microbiol. 17, 1742–1751 (2015).

19. LaCourse, K. D. et al. Conditional toxicity and synergy drive diversity among antibacterial effectors. Nat. Microbiol. 3, 440–446 (2018).

20. Ting, S. Y. et al. Targeted Depletion of bacteria from mixed populations by programmable adhesion with antagonistic competitor cells. Cell Host Microbe 28, 313-321.e6 (2020).

21. Hoff, j. et al. Vibrio natriegens?: an ultrafast-growing marine bacterium as emerging synthetic biology chassis. Environ. Microbiol. 1462–2920.15128 (2020) doi:10.1111/1462-2920.15128.

22. Weinstock, M. T., Hesek, E. D., Wilson, C. M. & Gibson, D. G. Vibrio natriegens as a fast-growing host for molecular biology. Nat. Methods 13, 849–851 (2016).

23. Lee, H. H. et al. Vibrio natriegens, a new genomic powerhouse. bioRxiv 058487 (2016) doi:10.1101/058487.

24. Dar, Y., Salomon, D. & Bosis, E. The antibacterial and anti-eukaryotic Type VI secretion system MIX-effector repertoire in Vibrionaceae. Mar. Drugs 16, 433 (2018).

25. Salomon, D., Gonzalez, H., Updegraff, B. L. & Orth, K. Vibrio parahaemolyticus Type VI secretion system 1 Is activated in marine conditions to target bacteria, and is differentially regulated from system 2. PLoS One 8, e61086 (2013).

26. Salomon, D. et al. Marker for Type VI secretion system effectors. Proc. Natl. Acad. Sci. 111, 9271–9276 (2014).

27. Salomon, D., Klimko, J. A. & Orth, K. H-NS regulates the Vibrio parahaemolyticus Type VI secretion system 1. Microbiol. (United Kingdom) 160, 1867–1873 (2014).

28. Jana, B., Fridman, C. M., Bosis, E. & Salomon, D. A modular effector with a DNase domain and a marker for T6SS substrates. Nat. Commun. 10, 3595 (2019).

29. Fridman, C. M., Keppel, K., Gerlic, M., Bosis, E. & Salomon, D. A comparative genomics methodology reveals a widespread family of membrane-disrupting T6SS effectors. Nat. Commun. 11, 1085 (2020).

30. Salomon, D. et al. Type VI secretion system toxins horizontally shared between marine bacteria. PLoS Pathog. 11, 1–20 (2015).

31. Li, P. et al. Acute aepatopancreatic necrosis disease-causing Vibrio parahaemolyticus strains maintain an antibacterial Type VI secretion system with versatile effector repertoires. Appl. Environ. Microbiol. 83, e00737–17 (2017).

32. Metzger, L. C., Matthey, N., Stoudmann, C., Collas, E. J. & Blokesch, M. Ecological implications of gene regulation by TfoX and TfoY among diverse Vibrio species. Environ. Microbiol. 21, 2231–2247 (2019).

33. Ben-Yaakov, R. & Salomon, D. The regulatory network of Vibrio parahaemolyticus Type VI secretion system 1. Environ. Microbiol. 21, 2248–2260 (2019).

34. Guzman, L. M., Belin, D., Carson, M. J. & Beckwith, J. Tight regulation, modulation, and high-level expression by vectors containing the arabinose P(BAD) promoter. J. Bacteriol. 177, 4121–4130 (1995).

35. Austin, B. Taxonomy of bacterial fish pathogens. Veterinary Research vol. 42 1–13 (2011).

36. Baker-Austin, C. et al. Vibrio spp. infections. Nat. Rev. Dis. Prim. 4, 1–19 (2018).

37. Ponce-Palafox, J., Martinez-Palacios, C. A. & Ross, L. G. The effects of salinity and temperature on the growth and survival rates of juvenile white shrimp, Penaeus vannamei, Boone, 1931. Aquaculture 157, 107–115 (1997).

38. Wyban, J., Walsh, W. A. & Godin, D. M. Temperature effects on growth, feeding rate and feed conversion of the Pacific white shrimp (Penaeus vannamei). Aquaculture 138, 267–279 (1995).

39. Banerjee, S., Petronella, N., Chew Leung, C. & Farber, J. Draft genome sequences of four Vibrio parahaemolyticus isolates from clinical cases in Canada. Genome Announc. 3, (2015).

40. Boyer, F., Fichant, G., Berthod, J., Vandenbrouck, Y. & Attree, I. Dissecting the bacterial Type VI secretion system by a genome wide in silico analysis: What can be learned from available microbial genomic resources? BMC Genomics 10, (2009).

41. Payne, W. J. Studies on bacterial utilization of uronic acids. III. Induction of oxidative enzymes in a marine isolate. J. Bacteriol. 76, 301–307 (1958).

42. Hersch, S. J. et al. Envelope stress responses defend against Type six secretion system attacks independently of immunity proteins. Nat. Microbiol. 5, 706–714 (2020).

43. Le, N. H. et al. Peptidoglycan editing provides immunity to Acinetobacter baumannii during bacterial warfare. Sci. Adv. 6, eabb5614 (2020).

44. Crisan, C. V. et al. Glucose confers protection to Escherichia coli against contact killing by Vibrio cholerae. Sci. Rep. 11, 2935 (2021).

45. Lin, H. H., Filloux, A. & Lai, E. M. Role of recipient susceptibility factors during contact-dependent interbacterial competition. Frontiers in Microbiology vol. 11 2768 (2020).

46. Gibson, D. G. et al. Enzymatic assembly of DNA molecules up to several hundred kilobases. Nat. Methods 6, 343–345 (2009).

47. Dunn, A. K., Millikan, D. S., Adin, D. M., Bose, J. L. & Stabb, E. V. New rfp- and pES213-derived tools for analyzing symbiotic Vibrio fischeri reveal patterns of infection and lux expression in situ. Appl. Environ. Microbiol. 72, 802–810 (2006).

48. O’Toole, R., Milton, D. L. & Wolf-Watz, H. Chemotactic motility is required for invasion of the host by the fish pathogen Vibrio anguillarum. Mol. Microbiol. 19, 625–637 (1996).

49. Bensadoun, A. & Weinstein, D. Assay of proteins in the presence of interfering materials. Anal. Biochem. 70, 241–250 (1976).

50. Untergasser, A. et al. Primer3-new capabilities and interfaces. Nucleic Acids Res. 40, e115–e115 (2012).

51. Schindelin, J. et al. Fiji: an open-source platform for biological-image analysis. Nat. Methods 9, 676–682 (2012).

