## Supplemental Data for "Engineering a customizable antibacterial T6SS-based platform in *Vibrio natriegens*"

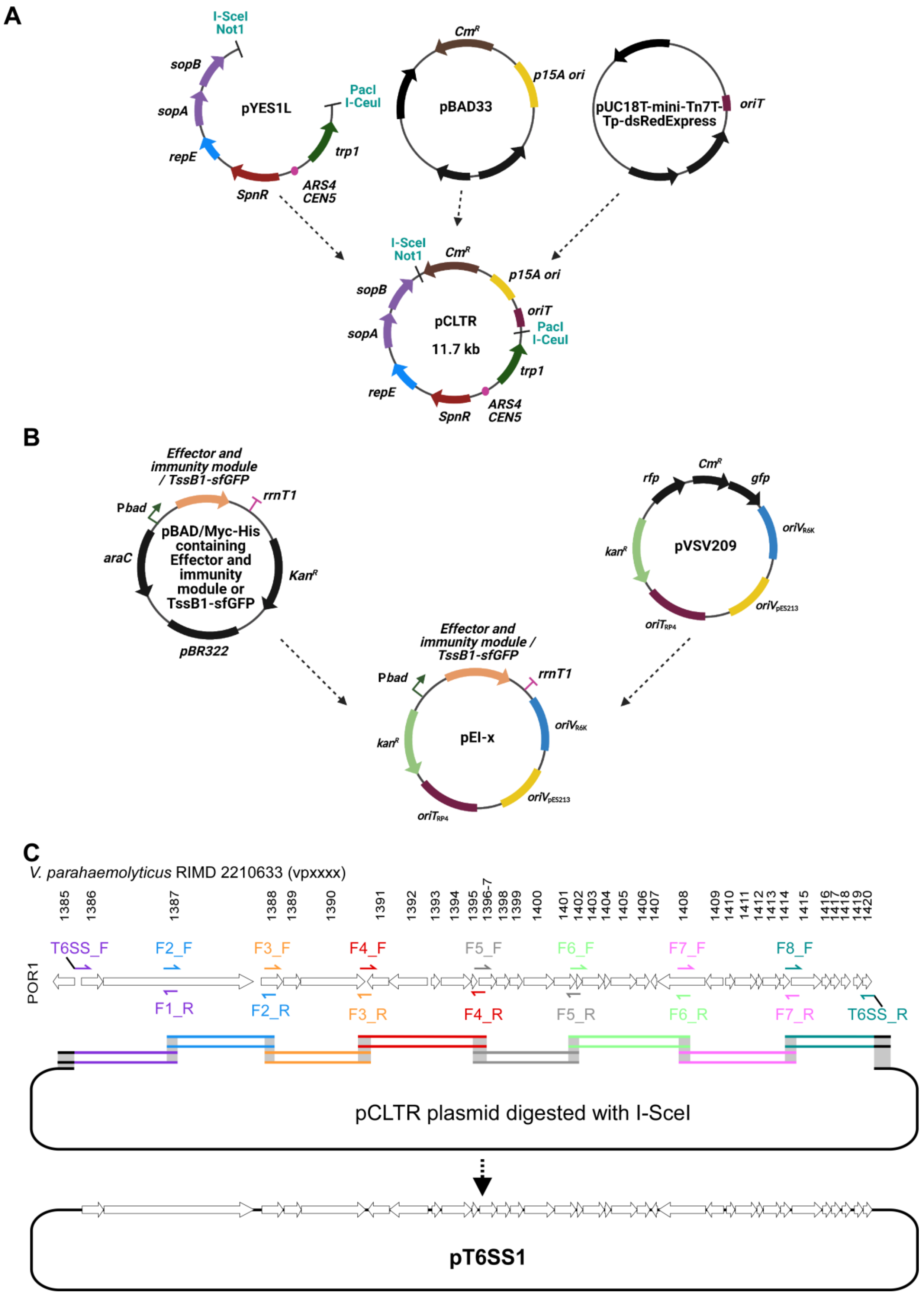
A
I-SceI
Not1
sopB
sopA
repE
SpnR
ARS4
CEN5
trp1
PacI
I-CeuI
pYES1L
CmR
p15A ori
pBAD33
oriT
pUC18T-mini-Tn7T-Tp-dsRedExpress
pCLTR
11.7 kb
B
Effector and immunity module / TssB1-sfGFP
rrnT1
Pbad
araC
pBAD/Myc-His containing Effector and immunity module or TssB1-sfGFP
KanR
pBR322
rfp
CmR
gfp
oriVR6K
kanR
pVSV209
oriTRP4
oriVpES213
pEI-x
C
V. parahaemolyticus RIMD 2210633 (vpxxxx)
1385 1386 1387 1388 1389 1390 1391 1392 1393 1394 1395 1396-7 1398 1399 1400 1401 1402 1403 1404 1405 1406 1407 1408 1409 1410 1411 1412 1413 1414 1415 1416 1417 1418 1419 1420
POR1
T6SS_F
F2_F
F3_F
F4_F
F5_F
F6_F
F7_F
F8_F
F1_R
F2_R
F3_R
F4_R
F5_R
F6_R
F7_R
T6SS_R
pCLTR plasmid digested with I-SceI
pT6SS1

**D**

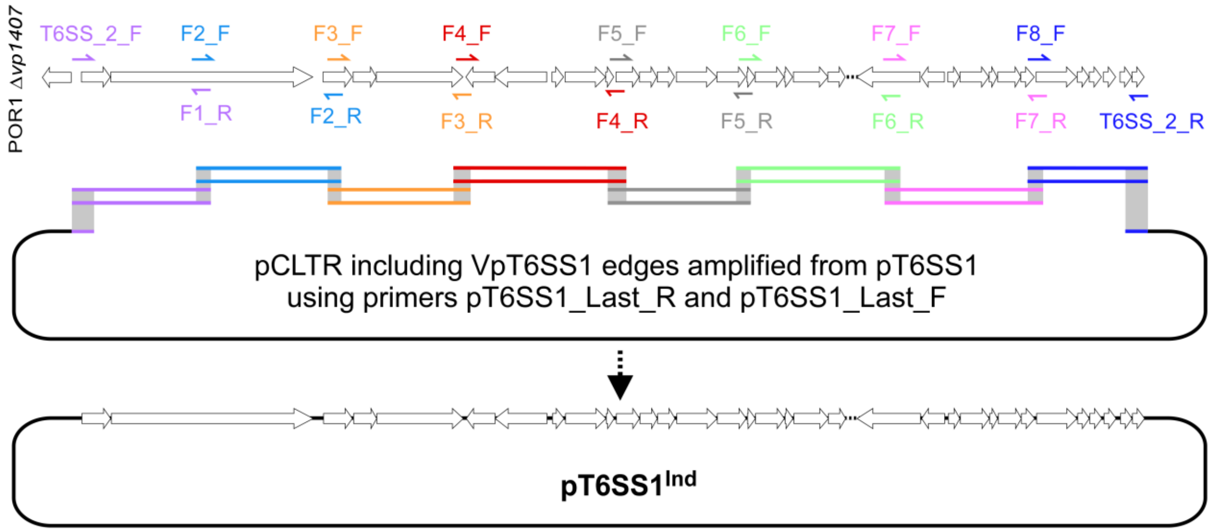
POR1 Δvp1407
T6SS_2_F
F2_F
F3_F
F4_F
F5_F
F6_F
F7_F
F8_F
F1_R
F2_R
F3_R
F4_R
F5_R
F6_R
F7_R
T6SS_2_R
pCLTR including VpT6SS1 edges amplified from pT6SS1
using primers pT6SS1_Last_R and pT6SS1_Last_F
pT6SS1Ind

**E**

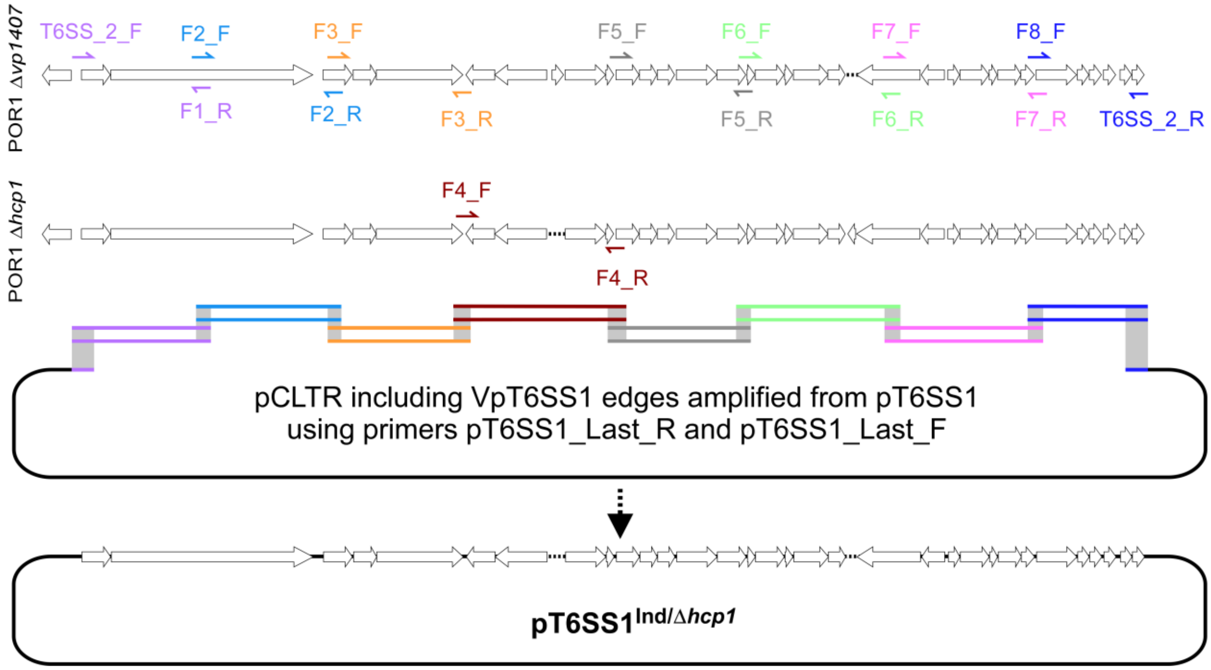
POR1 Δvp1407
T6SS_2_F
F2_F
F3_F
F5_F
F6_F
F7_F
F8_F
F1_R
F2_R
F3_R
F5_R
F6_R
F7_R
T6SS_2_R
POR1 Δhcp1
F4_F
F4_R
pCLTR including VpT6SS1 edges amplified from pT6SS1
using primers pT6SS1_Last_R and pT6SS1_Last_F
pT6SS1Ind/Δhcp1

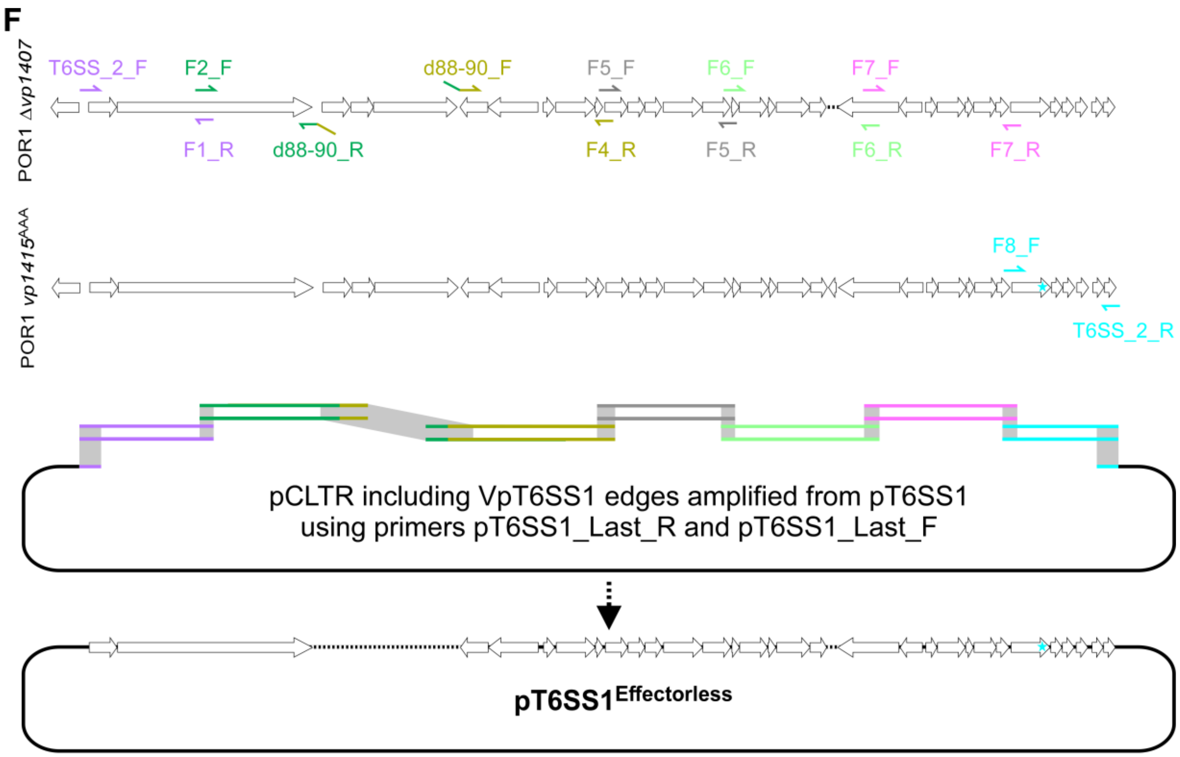
F
POR1 Δvp1407
T6SS_2_F
F2_F
d88-90_F
F5_F
F6_F
F7_F
F1_R
d88-90_R
F4_R
F5_R
F6_R
F7_R
POR1 vp1415AAA
F8_F
T6SS_2_R
pCLTR including VpT6SS1 edges amplified from pT6SS1
using primers pT6SS1_Last_R and pT6SS1_Last_F
pT6SS1Effectorless

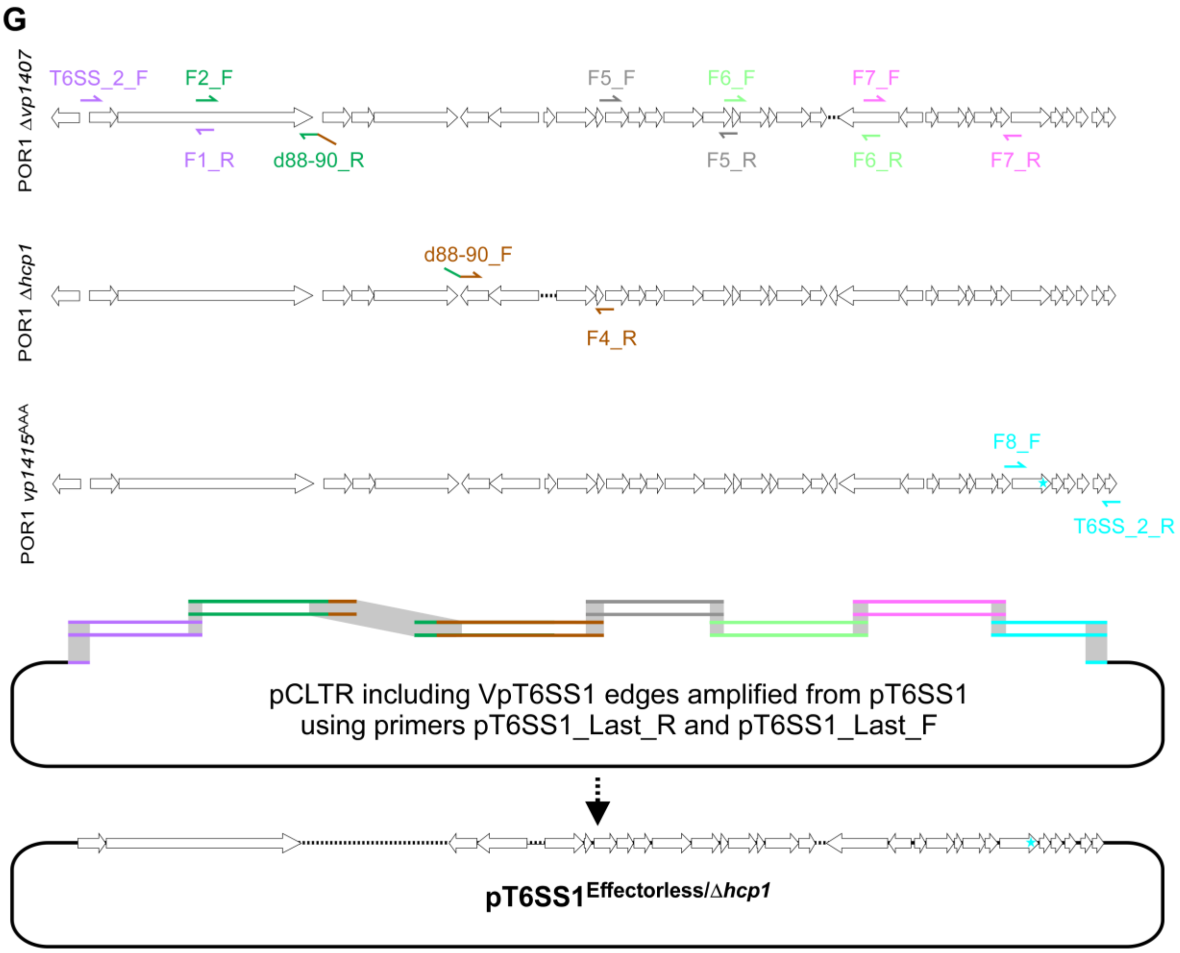
G
POR1 Δvp1407
T6SS_2_F
F2_F
F5_F
F6_F
F7_F
F1_R
d88-90_R
F5_R
F6_R
F7_R
POR1 Δhcp1
d88-90_F
F4_R
POR1 vp1415AAA
F8_F
T6SS_2_R
pCLTR including VpT6SS1 edges amplified from pT6SS1
using primers pT6SS1_Last_R and pT6SS1_Last_F
pT6SS1Effectorless/Δhcp1

H

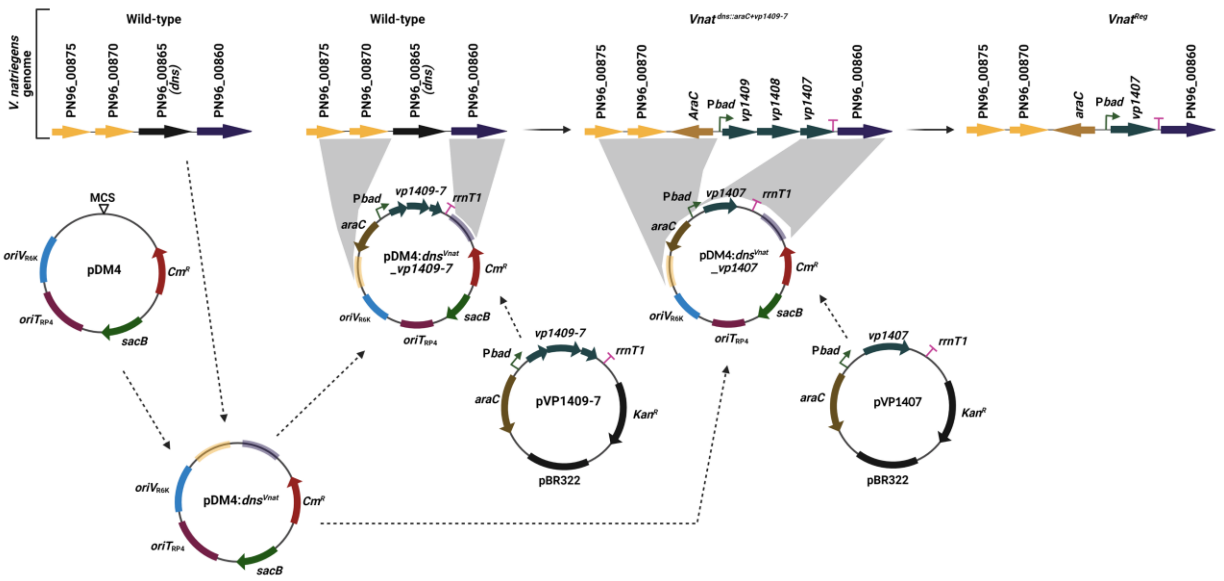


**Supplementary Figure S1. Schematic representation of plasmid and strain construction. A-B)** Construction of pCLTR (A) and pVSV209-derived (pEI-x) (B) plasmids. Plasmid names are denoted in the center of each circle. Genes are denoted as arrows in the direction of transcription. Bacterial origins of replication or conjugative transfer are denoted as rectangles. Yeast origin of replication is denoted as small circles. Notable restriction sites are denoted in turquoise. Sequence components that were amplified and used to construct downstream plasmids are colored. **C-G)** Construction of pT6SS1 and its derivatives. VpT6SS1 genes are denoted as white arrows in the direction of transcription. Locus numbers in *V. parahaemolyticus* RIMD 2210633 (vpxxxx) are denoted above the cluster shown in C. Primers used to amplify VpT6SS1 fragments are denoted as one-sided arrows and are color coded with their names. Colored lines denote amplified dsDNA fragments, and gray rectangles denote identical sequences shared between the amplified fragments and with I-SceI-digested or PCR amplified pCLTR (black). Dashed lines denote regions in the DNA from which genes were deleted. Replacing AAA with AHH in *vp1415* is denoted by a cyan star. **H)** Construction of *Vnat*$^{\text{Reg}}$. Plasmid names are denoted in the center of each circle. Genes are denoted as arrows in the direction of transcription. Bacterial origins of replication or conjugative transfer are denoted as solid rectangles. Sequence components that were amplified and used to construct downstream plasmids are colored. Dashed arrows denote components used to construct the destination plasmids. Unbroken, narrow black arrows denote the progression of *V. natriegens* genome engineering. Gray shapes denote identical sequences.

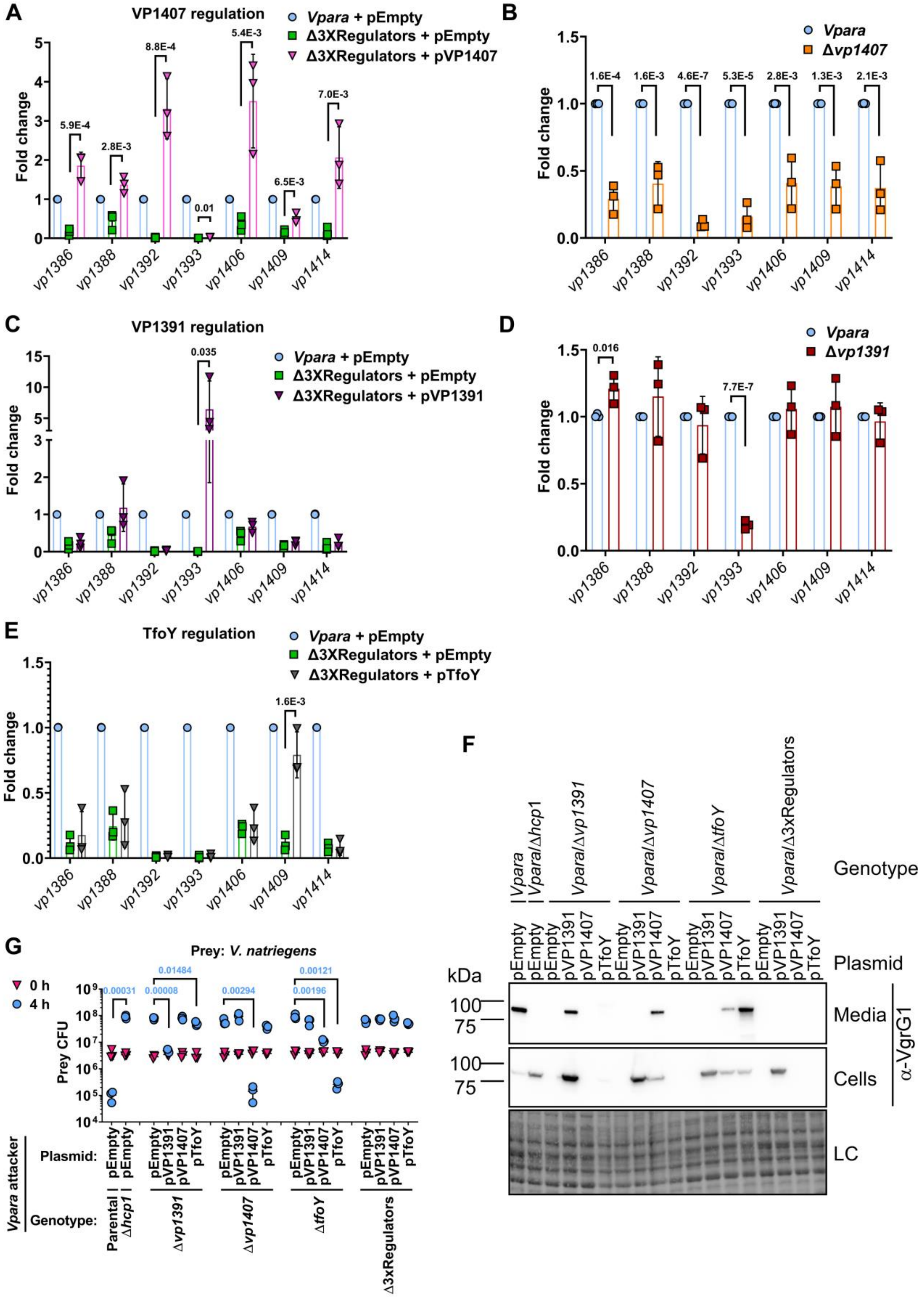
A
VP1407 regulation
Vpara + pEmpty
Δ3XRegulators + pEmpty
Δ3XRegulators + pVP1407
Fold change
5.9E-4
2.8E-3
8.8E-4
0.01
5.4E-3
6.5E-3
7.0E-3
vp1386 vp1388 vp1392 vp1393 vp1406 vp1409 vp1414
B
Vpara
Δvp1407
1.6E-4 1.6E-3 4.6E-7 5.3E-5 2.8E-3 1.3E-3 2.1E-3
Fold change
vp1386 vp1388 vp1392 vp1393 vp1406 vp1409 vp1414
C
VP1391 regulation
Vpara + pEmpty
Δ3XRegulators + pEmpty
Δ3XRegulators + pVP1391
0.035
Fold change
vp1386 vp1388 vp1392 vp1393 vp1406 vp1409 vp1414
D
Vpara
Δvp1391
0.016
7.7E-7
Fold change
vp1386 vp1388 vp1392 vp1393 vp1406 vp1409 vp1414
E
TfoY regulation
Vpara + pEmpty
Δ3XRegulators + pEmpty
Δ3XRegulators + pTfoY
1.6E-3
Fold change
vp1386 vp1388 vp1392 vp1393 vp1406 vp1409 vp1414
F
Vpara
Vpara/Δhcp1
Vpara/Δvp1391
Vpara/Δvp1407
Vpara/ΔtfoY
Vpara/Δ3xRegulators
Genotype
pEmpty pEmpty pEmpty pVP1391 pVP1407 pTfoY pEmpty pVP1391 pVP1407 pTfoY pEmpty pVP1391 pVP1407 pTfoY pEmpty pVP1391 pVP1407 pTfoY
Plasmid
kDa
100
75
Media
Cells
α-VgrG1
LC
G
Prey: V. natriegens
0 h
4 h
0.00031 0.00008 0.01484 0.00294 0.00196 0.00121
Prey CFU
Vpara attacker
Plasmid:
Genotype:
Parental
Δhcp1
Δvp1391
Δvp1407
ΔtfoY
Δ3xRegulators

**Supplementary Figure S2. The effects of VpT6SS1 regulators on cluster expression and activity in *V. parahaemolyticus*. A-E)** Quantitative real-time PCR showing the fold change in the expression of the indicated VpT6SS1 genes under VpT6SS1-inducing conditions. The values were generated by comparing the expression of genes in the indicated strains (Δ3xRegulators is Δ*vp1391*/Δ*vp1407*/Δ*tfoY*) to the parental *V. parahaemolyticus* POR1 strain containing an empty plasmid (*Vpara* + pEmpty in A, C, and E) or no plasmid (*Vpara* in B and D), using the ΔΔCt method. In A, C, and E, mutant strains harbor the indicated arabinose-inducible expression vectors for the regulators VP1407 (pVP1407; in A), VP1391 (pVP1391; in C), or TfoY (pTfoY; in E), or an empty plasmid (pEmpty). 16s rRNA was used as a reference gene. Data are shown as the mean ± SD of results obtained in three independent experiments. **F)** Expression (cells) and secretion (media) of VgrG1 from *V. parahaemolyticus* POR1 (*Vpara*; T6SS1$^{+}$), its T6SS1$^{-}$ mutant (Δ*hcp1*), and from single (Δ*vp1391*, Δ*vp1407*, and Δ*tfoY*) or triple (Δ3xRegulators) regulator deletion strains carrying the indicated arabinose-inducible plasmids. Samples were treated with 20 µM phenamil to activate surface sensing in media containing 3% NaCl at 30 °C. Loading control (LC) is shown for total protein lysates. **G)** Viability counts of *V. natriegens* prey before (0 h) and after (4 h) co-incubation with the indicated *V. parahaemolyticus* POR1 (*Vpara*) attackers as described in F, on media containing 3% NaCl and 0.1% arabinose at 30 °C. Data are shown as the mean ± SD. Statistical significance between samples by an unpaired, one (A-E), or two (G)-tailed Student's *t*-test are denoted above. In G, statistical significance is denoted for samples at the 4 h timepoint. A significant difference was considered as $P < 0.05$.

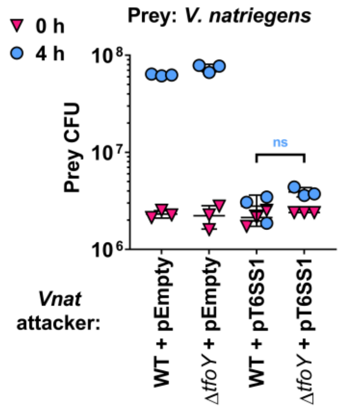


**Supplementary Figure S3. VpT6SS1 is not regulated by TfoY in *V. natriegens*.** Viability counts of *V. natriegens* prey before (0 h) and after (4 h) co-incubation with the indicated *V. natriegens* attackers containing the indicated plasmids on media containing 3% NaCl at 30 °C. Data are shown as the mean ± SD. Statistical significance between samples at the 4 h timepoint by an unpaired, two-tailed Student's *t*-test are denoted above. A significant difference was considered as $P < 0.05$. WT, wild-type; ns, not significant.

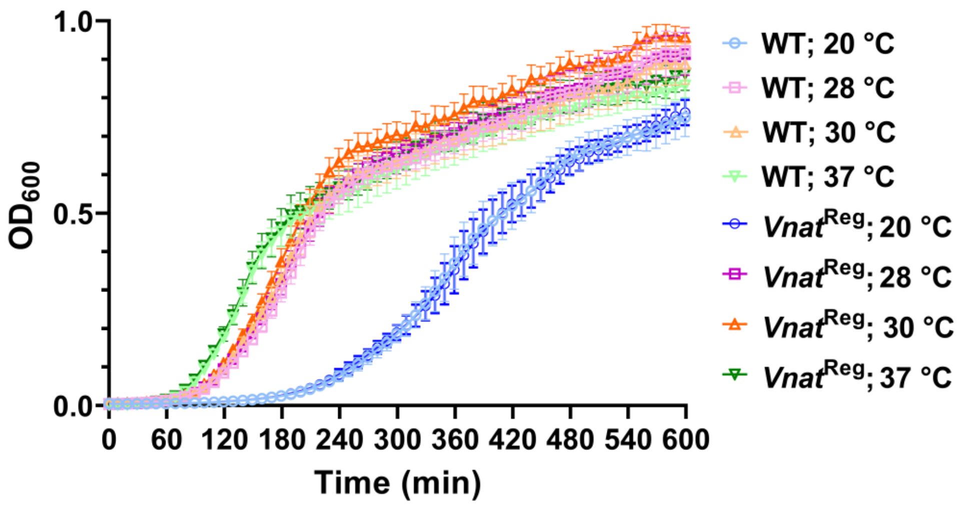


**Supplementary Figure S4. *V. natriegens* growth at different temperatures.** The growth of *V. natriegens* wild-type (WT) and the *Vnat*$^{Reg}$ derivative in media containing 3% NaCl at the indicated temperatures, as measured by $OD_{600}$ readings. Data are shown as the mean ± SD of 16 datapoints collected as technical quadruplicates in 4 independent experiments.

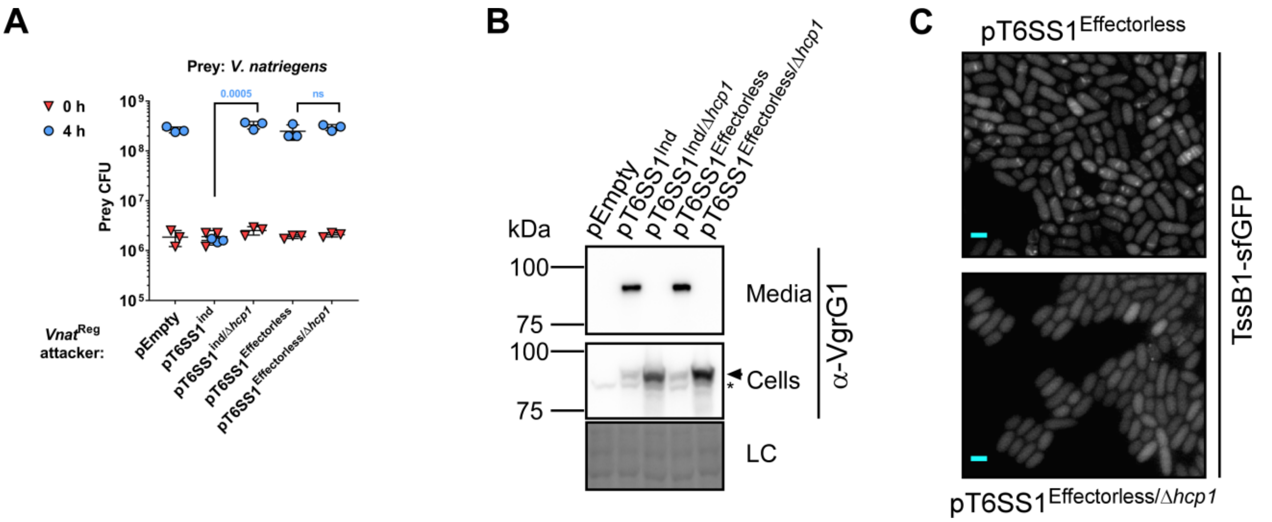


**Supplementary Figure S5. Constructing an effectorless VpT6SS1-based platform.** **A)** Viability counts of *V. natriegens* prey before (0 h) and after (4 h) co-incubation with *Vnat*$^{Reg}$ attackers harboring the indicated plasmids, on media containing 3% NaCl and 0.1% arabinose at 28 °C. Data are shown as the mean ± SD. Statistical significance between samples at the 4 h timepoint by an unpaired, two-tailed Student's *t*-test are denoted above. A significant difference was considered as $P < 0.05$. **B)** Expression (cells) and secretion (media) of VgrG1 from *Vnat*$^{Reg}$ harboring the indicated plasmids. Samples were grown in media containing 3% NaCl and 0.1% arabinose at 28 °C. Loading control (LC) is shown for total protein lysates. An arrow denotes bands corresponding to VgrG1. An asterisk denotes non-specific bands. **C)** Fluorescence microscopy of VpT6SS1 assembly in *Vnat*$^{Reg}$ containing the indicated pT6SS1 derivative plasmids and pTssB1-sfGFP for the arabinose-inducible expression of TssB1-sfGFP fusion. Representative images of TssB1-sfGFP signal are shown. Bar = 2 µm.

### Supplementary Table S1. Bacterial strains used in this study.

| Strain name | Genotype | Comments | Source |
|---|---|---|---|
| *Vibrio parahaemolyticus* RIMD 2210633 | Wild type | Used as a template for PCR amplifications | [1] |
| *Vibrio parahaemolyticus* RIMD 2210633 Δ*hcp1* | Δ*vp1393* | RIMD 2210633 derivative. Used as prey in bacterial competition assays | [2] |
| *Vibrio parahaemolyticus* POR1 | Δ*tdhAS* | RIMD 2210633 derivative. Used as an attacker in competition assays, in secretion assays, and as a template for PCR amplifications | [3] |
| *Vibrio parahaemolyticus* POR1 Δ*hcp1* | Δ*tdhAS*/Δ*vp1393* | RIMD 2210633 derivative. Used as an attacker in competition assays, in secretion assays, and as a template for PCR amplifications | [4] |
| *Vibrio parahaemolyticus* POR1 Δ*vp1391* | Δ*tdhAS*/Δ*vp1391* | RIMD 2210633 derivative. Used as an attacker in competition assays, in secretion assays, and for quantitative RT- PCR | [4] |
| *Vibrio parahaemolyticus* POR1 Δ*vp1407* | Δ*tdhAS*/Δ*vp1407* | RIMD 2210633 derivative. Used as an attacker in competition assays, in secretion assays, as a template for PCR amplifications, and for quantitative RT- PCR | [4] |
| *Vibrio parahaemolyticus* POR1 Δ*tfoY* | Δ*tdhAS*/Δ*vp1028* | RIMD 2210633 derivative. Used as an attacker in competition assays, in secretion assays, and for quantitative RT- PCR | [5] |
| *Vibrio parahaemolyticus* POR1 Δ3xRegulators | Δ*tdhAS*/Δ*vp1391*/Δ*vp1407*/Δ*vp1028* | RIMD 2210633 derivative. Used as an attacker in competition assays, in secretion assays, and for quantitative RT- PCR | This study |
| *Vibrio parahaemolyticus* POR1 *vp1415*$^{AAA}$ | Δ*tdhAS*/*vp1415*$^{AAA}$ | RIMD 2210633 derivative. Used as a template for PCR amplifications. Codons for histidines 563-4 in VP1415 were replaced by codons for alanines | This study |
| *Vibrio parahaemolyticus* 12-297/B Δ*hcp1* | Δ*b5c30_rs15290* | 12-297/B derivative. Used as prey in competition assays | [2] |

| | | | |
|---|---|---|---|
| *Vibrio parahaemolyticus* 04.2548 | Wild type | Used to prepare derivative strains | Obtained from Swapan Banerjee, [6] |
| *Vibrio parahaemolyticus* 04.2548 Δ*hcp1* | Δ*ba740_rs16850* | 04.2548 derivative. Used as prey in competition assays | This study |
| *Vibrio natriegens* ATCC 14048 | Wild type | Used as an attacker and prey in competition assays, as a template for PCR amplifications, and to prepare derivative strains | ATCC collection |
| *Vibrio natriegens* ATCC 14048 Δ*tfoY* | Δ*m272_rs24650* | ATCC 14048 derivative. Used as an attacker in competition assays | This study |
| *Vnat*$^{dns::araC+vp1409\text{-}7}$ | The endogenous *dns* gene (*pn96_00865*) replaced by a cassette containing *araC* and P*bad*-controlled *V. parahaemolyticus vp1409-7* | ATCC 14048 derivative. Used to prepare derivative strains | This study |
| *Vnat*$^{\mathrm{Reg}}$ | The endogenous *dns* gene (*pn96_00865*) replaced by a cassette containing *araC* and P*bad*-controlled *V. parahaemolyticus vp1407* | ATCC 14048 derivative. Used as an attacker in bacterial competition | This study |
| *Vibrio alginolyticus* 12G01 Δ*hcp1*/Δ*hcp2* | Δ*v12g01_01540*/Δ*v12g01_07583* | 12G01 derivative. Used as prey in competition assays | [7] |
| *Vibrio vulnificus* CMCP6 | Rifampicin-resistant parental strain | Used as prey in competition assays | Obtained from Karla Satchell |
| *Aeromonas jandaei* DSM 7311 | Wild type (natural resistance to ampicillin) | Used to prepare derivative strains | DSMZ collection |
| *Aeromonas jandaei* DSM 7311 Δ*tssB* | Δ*bn1126_rs13720* | DSM 7311 derivative. Used as prey in competition assays | This study |
| *Salmonella enterica* serover choleraesuis strain SC-B67 | Ampicillin, kanamycin, streptomycin and chloramphenicol-resistant strain | Used as prey in competition assays | Obtained from Ohad Gal-Mor |
| *Escherichia coli* DH5α | K-12 derivative laboratory strain | Used as prey in competition assays | Lab stocks |

| | | | |
|---|---|---|---|
| *Escherichia coli* DH5α (λ pir) | K-12 derivative laboratory strain containing λ pir | Used for plasmid maintenance and cloning, and as prey in competition assays | Obtained from Eric V. Stabb |

**Supplementary Table S2. Plasmids used in this study.**

| Plasmid name | Description | Purpose | Source |
|---|---|---|---|
| pYES1L | *E. coli*-yeast shuttle vector | Used to construct pCLTR | Novagen |
| pBAD33 | A pBAD series expression plasmid encoding the $Cm^R$ gene and the *p15A* origin of replication | $Cm^R$ and *p15A ori* were amplified to construct pCLTR | Addgene |
| pUC18T-mini-Tn7T-Tp-dsRedExpress | TpR mini-Tn7 vector, mobilizable, for RFP tagging bacteria | *oriT* was amplified to construct pCLTR | Addgene |
| sfGFP-N1 | Eukaryotic expression plasmid encoding superfolder GFP (sfGFP) | Used as a template to amplify *sfgfp* | Addgene |
| pEVS170 | A mini-Tn5 delivery vector; $Erm^R$, $Kan^R$ | Used to construct pTnp1222 | [8] |
| pTnp1222 | $Erm^R$-containing plasmid; constructed by first integrating pEVS170 into the genome of *V. parahaemolyticus* POR1 at a random location, followed by digestion of the genomic DNA with HhaI and self-ligation of the products | Used to selectively grow *E. coli* prey on erythromycin-containing media when 5 different prey strains were mixed together during bacterial competition | This study |
| pBAD18 | Bacterial expression vector with Gentamycin selection | Used to selectively grow *E. coli*, *V. natriegens*, *V. parahaemolyticus*, *V. alginolyticus*, and *A. jandaei* prey on media containing gentamicin during bacterial competition | Addgene |
| pBAD/*Myc*-His | Arabinose-inducible pBAD/*Myc*-His plasmid harboring a $Kan^R$ cassette | Used for cloning and arabinose-inducible expression of various genes in *Vibrio* | [4] |
| pVP1391 | pBAD/*Myc*-His containing the CDS of *vp1391* in the MCS, in-frame with the C-terminal *Myc*-His tag | Used for arabinose-inducible expression of VP1391 | This study |

| | | | |
|---|---|---|---|
| pVP1407 | pBAD/*Myc*-His containing the CDS of *vp1407* in the MCS, out-of-frame with the C-terminal *Myc*-His tag | Used for arabinose-inducible expression of VP1407 | This study |
| pTfoY | pBAD/*Myc*-His containing the CDS of *vp1028* in the MCS, in-frame with the C-terminal *Myc*-His tag | Used for arabinose-inducible expression of TfoY | [5] |
| pVP1409-7 | pBAD/*Myc*-His containing the CDS of *vp1409-7* in the MCS, out-of-frame with the C-terminal *Myc*-His tag | Used to place *vp1409-7* under a P*bad* promoter | This study |
| pBAD/*Myc*-His:PoNe/i$^{Vp\ 12\text{-}297/B}$ | pBAD/*Myc*-His containing the *V. parahaemolyticus* 12-297/B effector and immunity module PoNe/i$^{Vp\ 12\text{-}297/B}$, consisting of *b5c30_rs14470-60*, in its MCS; the 3' gene is cloned in-frame with the C-terminal *Myc*-His tag of the plasmid | Used to place PoNe/i$^{Vp\ 12\text{-}297/B}$ under P*bad* control | [9] |
| pBAD/*Myc*-His:Tme/i1$^{Vp\ BB22OP}$ | pBAD/*Myc*-His containing the *V. parahaemolyticus* BB22OP effector and immunity module Tme/i1$^{Vp\ BB22OP}$, consisting of *vpbb_rs15030-35*, in its MCS; the 3' gene is cloned in-frame with the C-terminal *Myc*-His tag of the plasmid | Used to place Tme/i1$^{Vp\ BB22OP}$ under P*bad* control | [9] |
| pBAD/*Myc*-His:VPA1263-Vti2$^{Vp\ RIMD}$ | pBAD/*Myc*-His containing the *V. parahaemolyticus* RIMD 2210633 effector and immunity module VPA1263-Vti2$^{Vp\ RIMD}$ in its MCS; the 3' gene is cloned in-frame with the C- | Used to place VPA1263-Vti2$^{Vp\ RIMD}$ under P*bad* control | This study |

| | | | |
|---|---|---|---|
| | terminal *Myc*-His tag of the plasmid | | |
| pBAD/*Myc*-His:Va2265-0$^{Va\ 12G01}$ | pBAD/*Myc*-His containing the *V. alginolyticus* 12G01 effector and immunity module Va02265-0$^{Va\ 12G01}$, consisting of *v12g01_02265-0*, in its MCS; the 3' gene is cloned out-of-frame with the C-terminal *Myc*-His tag of the plasmid | Used to place Va02265-0$^{Va\ 12G01}$ under P*bad* control | [7] |
| pBAD/*Myc*-His:TssB1-sfGFP | pBAD/*Myc*-His containing the *V. parahaemolyticus* RIMD 2210633 *tssB1* (*vp1402*) fused to a linker (AAAGGG) and *sfgfp*; 3' is cloned in-frame with the C-terminal *Myc*-His tag of the plasmid | Used to fuse TssB1 with sfGFP under P*bad* control | This study |
| pDM4 | a $Cm^R$ and $oriV_{R6K}$-containing suicide vector | To generate deletions, insertions, and replacements in *Vibrio* genomes | [10] |
| pDM4:*vp1391* | pDM4 containing 1 kb upstream and 1 kb downstream of *vp1391* in its MCS | Used to generate *V. parahaemolyticus* POR1 Δ3x Regulators | [4] |
| pDM4:*vp1407* | pDM4 containing 1 kb upstream and 1 kb downstream of *vp1407* in its MCS | Used to generate *V. parahaemolyticus* POR1 Δ3x Regulators | [4] |
| pDM4:*tfoY* | pDM4 containing 1 kb upstream and 1 kb downstream of the *V. parahaemolyticus vp1028* in its MCS | Used to generate *V. parahaemolyticus* POR1 Δ3x Regulators | [5] |
| pDM4:*dns*$^{Vnat}$ | pDM4 containing 1 kb upstream and 1 kb downstream of *V. natriegens dns* in its MCS | Used as platform to introduce cassettes that will replace *dns* in the *V. natriegens* genome | This study |
| pDM4:*dns*$^{Vnat}$_*vp1409-7* | pDM4:*dns*$^{Vnat}$ into which the region spanning *araC* to the *rrnT1* terminator from | Used to replace the *V. natriegens dns* gene with a cassette including a constitutively expressed | This study |

| | | | |
|---|---|---|---|
| | pVP1409-7 was inserted between the upstream and the downstream regions of *dns* | *araC* and *vp1409-7* under P*bad* control | |
| pDM4:*dns*$^{Vnat}$*_vp1407* | pDM4:*dns*$^{Vnat}$ into which the region spanning *araC* to the *rrnT1* terminator from pVP1407 was inserted between the upstream and downstream regions of *dns* | Used to replace the *V. natriegens dns* gene with a cassette including a constitutively expressed *araC* and *vp1407* under P*bad* control | This study |
| pDM4:*tfoY*$^{Vnat}$ | pDM4 containing 1 kb upstream and 1 kb downstream of the *V. natrigens m272_rs24650* in its MCS | Used to delete *tfoY* in *V. natriegens* | This study |
| pDM4:*vp1415* | pDM4 containing a ~2.2 kb region encompassing 1.1 kb upstream and 1.1 kb downstream of the codons encoding histidines 563-4 in *V. parahaemolyticus vp1415* | Used as a template to replace the codons encoding histidines 563-4 in *V. parahaemolyticus vp1415* with alanines | This study |
| pDM4:*vp1415*$^{AAA}$ | pDM4 containing a ~2.2 kb region encompassing 1.1 kb upstream and 1.1 kb downstream of the codons encoding histidines 563-4 in *V. parahaemolyticus vp1415* in which these two codons were mutated to encode alanines | Used to replace VP1415 histidines 563-4 with alanine in the *V. parahaemolyticus* genome | This study |
| pDM4:*tssB*$^{Aj\ DSM7311}$ | pDM4 containing 1 kb upstream and 1 kb downstream of *A. jandaei bn1126_rs13720* | Used to delete *tssB* in *A. jandaei* | This study |
| pDM4:*hcp1*$^{Vp\ 04.2548}$ | pDM4 containing 1 kb upstream and 1 kb downstream of *V. parahaemolyticus* | Used to delete *hcp1* in *V. parahaemolyticus* 04.2548 | This study |

| | | | |
|---|---|---|---|
| | 04.2548 *ba740_rs16850* | | |
| pVSV209 | Mobilizable plasmid contains $OriV_{pES213}$, $OriV_{R6K}$, $Kan^R$, constitutive *rfp*, and a promoterless $Cm^R$-*gfp* reporter | Used to carry TssB1-sfGFP and effector and immunity modules in *V. natriegens* | [11] |
| pBJ209-araC | pVSV209 derivative in which the *rfp-gfp* cassette was replaced by the region spanning *araC* CDS to the *rnnT1* terminator from pBAD/*Myc*-His | Used as pEmpty for pVSV209-derived plasmids (pEI-x) | This study |
| pPoNe/i$^{Vp\ 12\text{-}297/B}$ | pVSV209 derivative in which the *rfp-gfp* cassette was replaced by the region spanning the *Pbad* promoter to the *rnnT1* terminator from pBAD/*Myc*-His:PoNe/i$^{Vp\ 12\text{-}297/B}$ | Used to express PoNe/i$^{Vp\ 12\text{-}297/B}$ in *V. natriegens* | This study |
| pTme/i1$^{Vp\ BB22OP}$ | pVSV209 derivative in which the *rfp-gfp* cassette was replaced by the region spanning the *Pbad* promoter to the *rnnT1* terminator from pBAD/*Myc*-His: Tme/i1$^{Vp\ BB22OP}$ | Used to express Tme/i1$^{Vp\ BB22OP}$ in *V. natriegens* | This study |
| pVPA1263-Vti2$^{Vp\ RIMD}$ | pVSV209 derivative in which the *rfp-gfp* cassette was replaced by the region spanning the *Pbad* promoter to the *rnnT1* terminator from pBAD/*Myc*-His: VPA1263-Vti2$^{Vp\ RIMD}$ | Used to express VPA1263-Vti2$^{Vp\ RIMD}$ in *V. natriegens* | This study |
| pVa2265-0$^{Va\ 12G01}$ | pVSV209 derivative in which the *rfp-gfp* cassette was replaced by the region spanning the *Pbad* promoter to the *rnnT1* terminator from | Used to express Va2265-0$^{Va\ 12G01}$ in *V. natriegens* | This study |

| | | | |
|---|---|---|---|
| | pBAD/*Myc*-His: Va2265-0$^{Va\ 12G01}$ | | |
| pVPA1263-Vti2+Va02265-0 | pVSV209 derivative in which the *rfp-gfp* cassette was replaced by the region spanning the *Pbad* promoter to the *rnnT1* terminator from pBAD/*Myc*-His: VPA1263-Vti2$^{Vp\ RIMD}$ followed by the same region from pBAD/*Myc*-His: Va2265-0$^{Va\ 12G01}$ | Used to express VPA1263-Vti2$^{Vp\ RIMD}$ and Va2265-0$^{Va\ 12G01}$ in *V. natriegens* together | This study |
| pTssB1-sfGFP | pVSV209 derivative in which the *rfp-gfp* cassette was replaced by the region spanning the *Pbad* promoter to the *rnnT1* terminator from pBAD/*Myc*-His:TssB1-sfGFP | Used to express TssB1-sfGFP in *V. natriegens* together | This study |
| pCLTR | *E. coli*-yeast-*Vibrio* shuttle vector; mobilizable; it contains $Spec^R$ and $Cm^R$ | Used to capture VpT6SS1 and its derivatives | This study |
| pT6SS1 | *vp1386-vp1420* (VpT6SS1) from *V. parahaemolyticus* in pCLTR | Used to transfer VpT6SS1 into *V. natriegens* | This study |
| pT6SS1$^{Ind}$ | pT6SS1 lacking *vp1407* | Used as an inducible derivative of pT6SS1 | This study |
| pT6SS1$^{Ind/\Delta hcp1}$ | pT6SS1$^{Ind}$ lacking *hcp1* | Used as a T6SS-inactive version of pT6SS1$^{Ind}$ | This study |
| pT6SS1$^{Effectorless}$ | pT6SS1$^{Ind}$ lacking *vp1388-90* and carrying a mutated form of *vp1415* (*vp1415*$^{AAA}$) | Used as an inducible derivative of pT6SS1 lacking endogenous active effector and immunity modules | This study |
| pT6SS1$^{Effectorless/\Delta hcp1}$ | pT6SS1$^{Effectorless}$ lacking *hcp1* | Used as a T6SS-inactive version of pT6SS1$^{Effectorless}$ | This study |

**Supplementary Table S3. Primers used in this study.**

| Primer name | Sequence (5' to 3') | Description |
|---|---|---|
| pCLTR_F1 | GCGCAGCGGCGGCCGCGCTGATACCGCCGCTTGCTTTCGAATTTCTGCCATTC | Amplify *p15A ori* and $Cm^R$ from pBAD33 to construct pCLTR |
| pCLTR_R1 | GCCGCAGCCGAACGACCGAGATAAGCTGTCAAACATGAGCAG | Amplify *p15A ori* and $Cm^R$ from pBAD33 to construct pCLTR |
| pCLTR_F2 | TGCTCATGTTTGACAGCTTATCTCGGTCGTTCGGCTGCGGCG | Amplify *oriT* from pUC18-mini-Tn7T-Tp-dsRedExpress to construct pCLTR |
| pCLTR_R2 | TCGCTCACTGACTTTAATTAACTGCGGCGAGGTATAGCGATTTTTTCGGTATATCC | Amplify *oriT* from pUC18-mini-Tn7T-Tp-dsRedExpress to construct pCLTR |
| T6SS_F | CCCTGTTTGCATTATGAATTAGTTACGCTAGGGGCGGCCGCAGTGAGCAGATAATGACTAAAAGATAC | Amplify a fragment of VpT6SS1 to construct pT6SS1 |
| F1_R | GCCTTTACGGTAGCGCTTGTTGATTTC | Amplify a fragment of VpT6SS1 to construct pT6SS1 and its derivatives |
| F2_F | TGTAACAGCGTATGTTACAGATAAAAAC | Amplify a fragment of VpT6SS1 to construct pT6SS1 and its derivatives |
| F2_R | TTTATTTATCTGGCTTCGTAACATTGC | Amplify a fragment of VpT6SS1 to construct pT6SS1 and its derivatives |
| F3_F | GAAGTATATTTGGCTTATAACAGTAAAC | Amplify a fragment of VpT6SS1 to construct pT6SS1 and its derivatives |
| F3_R | ATACGACATTTCTGCGGCAGGTTG | Amplify a fragment of VpT6SS1 to construct pT6SS1 and its derivatives |
| F4_F | CTTGGAAGCAGTATAACGAATCTCAAC | Amplify a fragment of VpT6SS1 to construct pT6SS1 and its derivatives |
| F4_R | GTTGTTAGCGCGCGATAGC | Amplify a fragment of VpT6SS1 to construct pT6SS1 and its derivatives |
| F5_F | CAGTGTTAACTAATTACATCGTG | Amplify a fragment of VpT6SS1 to construct pT6SS1 and its derivatives |
| F5_R | GACATAATTAGTTCCTTATCTAAC | Amplify a fragment of VpT6SS1 to construct pT6SS1 and its derivatives |

| | | |
|---|---|---|
| F6_F | GAAAAGTGGAAAGCAATTTATGAC | Amplify a fragment of VpT6SS1 to construct pT6SS1 and its derivatives |
| F6_R | CTACTTCCGCACTCAATTCTCACC | Amplify a fragment of VpT6SS1 to construct pT6SS1 and its derivatives |
| F7_F | AGTTGCTCGAACGACATCCATGTTG | Amplify a fragment of VpT6SS1 to construct pT6SS1 and its derivatives |
| F7_R | CCACTCTTCGCCAGTCTTATCCAG | Amplify a fragment of VpT6SS1 to construct pT6SS1 and its derivatives |
| F8_F | GCATTGCCTTACCACGCAGGTTAC | Amplify a fragment of VpT6SS1 to construct pT6SS1 and its derivatives |
| T6SS_R | GCGCTCGGTCGTTCGGGTTCTATATTACCCTGGCGGCCGCAAGTGGTACTTCACTTGTTCAGCCG | Amplify a fragment of VpT6SS1 to construct pT6SS1 |
| T6SS_2_R | AAGTGGTACTTCACTTGTTCAGCC | Amplify a fragment of VpT6SS1 to construct pT6SS1 derivatives |
| T6SS_2_F | AGTGAGCAGATAATGACTAAAAGATAC | Amplify a fragment of VpT6SS1 to construct pT6SS1 derivatives |
| d88-90_R | GTTGAGATTCGTTATACTGCTTCCAAGCCAACATCGTAAACATGGTTGCGG | Amplify a fragment of VpT6SS1 to construct pT6SS1 derivatives |
| d88-90_F | CCGCAACCATGTTTACGATGTTGGCTTGGAAGCAGTATAACGAATCTCAAC | Amplify a fragment of VpT6SS1 to construct pT6SS1 derivatives |
| pT6SS1_Last_R | ATATCTATCATTATTTAGTTAGTAAAAACAAAGGAATTGTC | Amplify the pT6SS1 backbone to construct pT6SS1 derivatives |
| pT6SS1_Last_F | AAAGTGCCGAAAGGCGCTTTTTCTTTTAATTACAATCC | Amplify the pT6SS1 backbone to construct pT6SS1 derivatives |
| pBADfix_GIB_F | AAGCTTGGGCCCGAACAAAAAC | Amplify the pBAD/*Myc*-His backbone for cloning using the Gibson assembly method |
| pBADfix_GIB_R | CATGGTTAATTCCTCCTGTTAG | Amplify the pBAD/*Myc*-His backbone for cloning using the Gibson assembly method |
| pBAD_vp1391_F | GCTAACAGGAGGAATTAACCATGCGTTCAGCTAACCATAGCG | Amplify *vp1391* to construct pVP1391 |
| pBAD_vp1391_R | TTTTGTTCGGGCCCAAGCTTACCTGCCGCAGAAATGTCGTATTTC | Amplify *vp1391* to construct pVP1391 |

| | | |
|---|---|---|
| pBAD_vp1407_F | GCTAACAGGAGGAATTAACCATGATTTCTAATATGACTTTACAAG | Amplify *vp1407* to construct pVP1407 |
| pBAD_vp1407_R_STOP | TTTTGTTCGGGCCCAAGCTTCTATCGACTGATCGGCTGATGAAG | Amplify *vp1407* to construct pVP1407 and pVP1409-7 |
| pBAD_VP1409_F | GCTAACAGGAGGAATTAACCATGGAACTTAACATTATGTTTTTCAG | Amplify *vp1409* to construct pVP1409-7 |
| VN_Dns_Up_F | CGCGAGCTCTTGCCCTTCTTAGTGATTGGGTC | Amplify the 1 kb upstream of *V. natriegens dns* (PN96_00865) to construct pDM4:*dns*$^{Vnat}$ |
| VN_Dns_Up_R | ATAAGAATGCGGCCGCAGTTAAAGTCTTTAAAAAGTATGAC | Amplify the 1 kb upstream of *V. natriegens dns* (PN96_00865) to construct pDM4:*dns*$^{Vnat}$ |
| VN_Dns_Dn_F | ATAAGAATGCGGCCGCCCAGCCGATAAATGGGAATGC | Amplify the 1 kb downstream of *V. natriegens dns* (PN96_00865) to construct pDM4:*dns*$^{Vnat}$ |
| VN_Dns_Dn_R | ACGCGTCGACGGCAAAGCTAGAGTCTTTCTTG | Amplify the 1 kb downstream of *V. natriegens dns* (PN96_00865) to construct pDM4:*dns*$^{Vnat}$ |
| Not1_Fix_F | ATAAGAATGCGGCCGCATGCATAATGTGCCTGTCAAATGG | Amplify the region spanning *araC* and *rrnT1* from pBAD/*Myc*-His derivatives to construct pDM4:*dns*$^{Vnat}$_*vp1407* or pDM4:*dns*$^{Vnat}$_*vp1409-7* |
| Not1_Fix_R | ATAAGAATGCGGCCGCTAGAAACGCAAAAAGGCCATCCG | Amplify the region spanning *araC* and *rrnT1* from pBAD/*Myc*-His derivatives to construct pDM4:*dns*$^{Vnat}$_*vp1407* or pDM4:*dns*$^{Vnat}$_*vp1409-7* |
| VN_TfoY_Up_F | CGCGAGCTCTATCGAATTGGCAGAGTACTTC | Amplify 1 kb upstream of *V. natriegens tfoY* (*m272_rs24650*) to construct pDM4:*tfoY*$^{Vnat}$ |
| VN_TfoY_Up_R | CGCGGATCCGATGTGTTATCTCGGTGTAATTATTATTACG | Amplify 1 kb upstream of *V. natriegens tfoY* (*m272_rs24650*) to construct pDM4:*tfoY*$^{Vnat}$ |
| VN_TfoY_Dn_F | CGCGGATCCTCTGTAGGTATAAACCTATCGTAC | Amplify 1 kb downstream of *V. natriegens tfoY* |

| | | |
|---|---|---|
| | | (*m272_rs24650*) to construct pDM4:*tfoY*$^{Vnat}$ |
| VN_TfoY_Dn_R | ACGCGTCGACAATGGTAGTTCCCCCTTCAATCCG | Amplify 1 kb downstream of *V. natriegens tfoY* (*m272_rs24650*) to construct pDM4:*tfoY*$^{Vnat}$ |
| VPA1263_F | TAACAGGAGGAATTAACCATGTCCTATAAAGTTCTTCCACTG | Amplify the *vpa1263*-*vti2* effector and immunity module from *V. parahaemolyticus* RIMD 2210633 to construct pBAD/*Myc*-His:VPA1263-Vti2$^{Vp\ RIMD}$ |
| Vit2_R | TTTTGTTCGGGCCCAAGCTTCTCTTTGAAGCAAGGTTCTC | Amplify the *vpa1263*-*vti2* effector and immunity module from *V. parahaemolyticus* RIMD 2210633 to construct pBAD/*Myc*-His:VPA1263-Vti2$^{Vp\ RIMD}$ |
| pVSV_F | AAACTCTTTTGTTTATTTTCCCCGGGAATTCCCATGTCAGCCG | Amplify the pVSV209 backbone for cloning using the Gibson assembly method |
| pVSV_R | GGCGGGAGTATGAAAAGGAGTGAGCTAACTCACATTAATTG | Amplify the pVSV209 backbone for cloning using the Gibson assembly method |
| pBADfix_R_MCS | AAAATAAACAAAAGAGTTTGTAGAAAC | Amplify the region between the P*bad* promoter and the *rnnT1* terminator from pBAD/*Myc*-His derivatives to clone into pVSV209 using the Gibson assembly method |
| pBADfix_F_MCS | CTTTTCATACTCCCGCCATTCAGAG | Amplify the region between the P*bad* promoter and the *rnnT1* terminator from pBAD/*Myc*-His derivatives to clone into pVSV209 using the Gibson assembly method |
| pDM4_Gib_VP1415_F | GTGGAATCCCGGGAGAGCTCAGAGATATAGAGCCAATCCAAAAG | Amplify the ~2.2 kb region flanking the *vp1415* codons encoding histidines 563-4 from *V. parahaemolyticus* POR1 to construct pDM4:*vp1415* |

| | | |
|---|---|---|
| pDM4_Gib_VP1415_R | TAGCGGAGTGTATATCAAGCTTTTAAATTTTCATTCCAAAACCAATG | Amplify the ~2.2 kb region flanking the *vp1415* codons encoding histidines 563-4 from *V. parahaemolyticus* POR1 to construct pDM4:*vp1415* |
| VP1415_HH_AA_F | GGAGCAGTGCAGGCTGCAGCTCTTATTCCTAAAAATGCATTC | Site-directed mutagenesis to replace histidines 563-4 of VP1415 with alanines |
| VP1415_HH_AA_R | CACTGCTCCCGCCCCGTGCCACCAAGGGTGC | Site-directed mutagenesis to replace histidines 563-4 of VP1415 with alanines |
| VP1386_F_RTPCR | AAGGCGTGGTGAACTTCAGT | Determine the expression of *vp1386* in quantitative RT-PCR |
| VP1386_R_RTPCR | CAAAGCTACTCTGCCCACCT | Determine the expression of *vp1386* in quantitative RT-PCR |
| VP1388_F_RTPCR | ACTGGGTTGCGTTGTCTTCT | Determine the expression of *vp1388* in quantitative RT-PCR |
| VP1388_R_RTPCR | CTGGACTTGGCTCAGAAGGG | Determine the expression of *vp1388* in quantitative RT-PCR |
| VP1392_F_RTPCR | ACCCTATTGCCGTTGGTGAG | Determine the expression of *vp1392* in quantitative RT-PCR |
| VP1392_R_RTPCR | ATGATCGGTGTTGGCGAGTT | Determine the expression of *vp1392* in quantitative RT-PCR |
| VP1393_F_RTPCR | AGCTGATTGATGCGCTATTGG | Determine the expression of *vp1393* in quantitative RT-PCR |
| VP1393_R_RTPCR | TCGTCGTTACCAGCTGTACC | Determine the expression of *vp1393* in quantitative RT-PCR |
| VP1406_F_RTPCR | CGGGCAGCAGACGATTCTAT | Determine the expression of *vp1406* in quantitative RT-PCR |
| VP1406_R_RTPCR | ACTCAATTCGCTCCTGGTGG | Determine the expression of *vp1406* in quantitative RT-PCR |
| VP1409_F_RTPCR | CCAGTCAAAGTGAGCCAGGT | Determine the expression of *vp1409* in quantitative RT-PCR |
| VP1409_R_RTPCR | GCAATGTTGGCAAAGACCGT | Determine the expression of *vp1409* in quantitative RT-PCR |

| | | |
|---|---|---|
| VP1414_F_RT-pcr | TTGGAGCCAGAACAGTTTGC | Determine the expression of *vp1414* in quantitative RT-PCR |
| VP1414_R_RT-pcr | TGCTAGCTTTTCGTTACCGC | Determine the expression of *vp1414* in quantitative RT-PCR |
| 16s rRNA_F RT-PCR | GACACGGTCCAGACTCCTAC | Determine the expression of 16s rRNA in quantitative RT-PCR |
| 16s rRNA_R RT-PCR | GGTGCTTCTTCTGTCGCTAAC | Determine the expression of 16s rRNA in quantitative RT-PCR |
| Aero_tssB_SalI_UP_F | CAAAGTCGACATGGTCTCGCCCTGAAAC | Amplify the 1 kb upstream of *A. jandaei* DSM 7311 *tssB* (*bn1126_rs13720*) to construct pDM4:*tssB*$^{Aj}_{DSM7311}$ |
| Aero_tssB_KpnI_UP_R_ | CAATGGTACCGCTCTAAATACCAAGTTAAGGG | Amplify the 1 kb upstream of *A. jandaei* DSM 7311 *tssB* (*bn1126_rs13720*) to construct pDM4:*tssB*$^{Aj}_{DSM7311}$ |
| Aero_tssB_KpnI_DN_F | CAATGGTACCTCGCCGGGTTGGATTGACTGAAC | Amplify the 1 kb downstream of *A. jandaei* DSM 7311 *tssB* (*bn1126_rs13720*) to construct pDM4:*tssB*$^{Aj}_{DSM7311}$ |
| Aero_tssB_XbaI_DN_R | CAAATCTAGAAGATCGTGGATGGCACCAC | Amplify the 1 kb downstream of *A. jandaei* DSM 7311 *tssB* (*bn1126_rs13720*) to construct pDM4:*tssB*$^{Aj}_{DSM7311}$ |
| TssB-link-GFP_F1 | GCTAACAGGAGGAATTAACCATGTCACGTGACGGCTCGGTG | Amplify *tssB1* (*vp1402*) from *V. parahaemolyticus* POR1 to construct pBAD/*Myc*-His:TssB1-sfGFP |
| TssB-link-sfGFP_R1 | ACCACCACCAGCAGCAGCCTCTTCTTTCGCGTCTTGGTC | Amplify *tssB1* (*vp1402*) from *V. parahaemolyticus* POR1 to construct pBAD/*Myc*-His:TssB1-sfGFP |
| TssB-link-sfGFP_F2 | GCTGCTGCTGGTGGTGGTATGGTGAGCAAGGGCGAGGAG | Amplify *sfgfp* from sfGFP-N1 plasmid to construct pBAD/*Myc*-His:TssB1-sfGFP |
| TssB-link-sfGFP_R2 | TTTTGTTCGGGCCCAAGCTTCTTGTACAGCTCGTCCATGCC | Amplify *sfgfp* from sfGFP-N1 plasmid to construct |

| | | |
|---|---|---|
| | | pBAD/*Myc*-His:TssB1-sfGFP |
| Hcp1_UP_F_SacI | CAAAGAGCTCCTGTCGTGAACTTGCTCAG | Amplify 1 kb upstream of *V. parahaemolyticus* 04.2548 *hcp1* (*ba740_rs16850*) to construct pDM4:$hcp1^{Vp}_{04.2548}$ |
| Hcp1_UP_R_BamHI | CAACGGATCCCGCTATTTCCTTTTCTAAAATCTG | Amplify 1 kb upstream of *V. parahaemolyticus* 04.2548 *hcp1* (*ba740_rs16850*) to construct pDM4:$hcp1^{Vp}_{04.2548}$ |
| Hcp1_DN_F_BamHI | CACCGGATCCTTGCTTTTTGCGTAAAGATTCAGG | Amplify 1 kb downstream of *V. parahaemolyticus* 04.2548 *hcp1* (*ba740_rs16850*) to construct pDM4:$hcp1^{Vp}_{04.2548}$ |
| Hcp1_DN_R_SalI | CAAAGTCGACCTGTAACCAGACGCCAAACG | Amplify 1 kb downstream of *V. parahaemolyticus* 04.2548 *hcp1* (*ba740_rs16850*) to construct pDM4:$hcp1^{Vp}_{04.2548}$ |

### SUPPLEMENTARY REFERENCES


1. Makino, K. *et al.* Genome sequence of *Vibrio parahaemolyticus*: a pathogenic mechanism distinct from that of V cholerae. *Lancet* **361**, 743–749 (2003).

2. Jana, B., Fridman, C. M., Bosis, E. & Salomon, D. A modular effector with a DNase domain and a marker for T6SS substrates. *Nat. Commun.* **10**, 3595 (2019).

3. Park, K.-S. *et al.* Functional characterization of two type III secretion systems of *Vibrio parahaemolyticus*. *Infect. Immun.* **72**, 6659–65 (2004).

4. Salomon, D., Gonzalez, H., Updegraff, B. L. & Orth, K. *Vibrio parahaemolyticus* Type VI secretion system 1 Is activated in marine conditions to target bacteria, and is differentially regulated from system 2. *PLoS One* **8**, e61086 (2013).

5. Ben-Yaakov, R. & Salomon, D. The regulatory network of *Vibrio parahaemolyticus* type VI secretion system 1. *Environ. Microbiol.* **21**, 2248–2260 (2019).

6. Banerjee, S., Petronella, N., Chew Leung, C. & Farber, J. Draft genome sequences of four *Vibrio parahaemolyticus* isolates from clinical cases in Canada. *Genome Announc.* **3**, (2015).

7. Salomon, D. *et al.* Type VI secretion system toxins horizontally shared between marine bacteria. *PLoS Pathog.* **11**, 1–20 (2015).

8. Lyell, N. L., Dunn, A. K., Bose, J. L., Vescovi, S. L. & Stabb, E. V. Effective mutagenesis of *Vibrio fischeri* by using hyperactive mini-Tn5 derivatives. *Appl. Environ. Microbiol.* **74**, 7059–63 (2008).

9. Fridman, C. M., Keppel, K., Gerlic, M., Bosis, E. & Salomon, D. A comparative genomics methodology reveals a widespread family of membrane-disrupting T6SS effectors. *Nat. Commun.* **11**, 1085 (2020).

10. O'Toole, R., Milton, D. L. & Wolf-Watz, H. Chemotactic motility is required for invasion of the host by the fish pathogen *Vibrio anguillarum*. *Mol. Microbiol.* **19**, 625–637 (1996).

11. Dunn, A. K., Millikan, D. S., Adin, D. M., Bose, J. L. & Stabb, E. V. New *rfp*- and pES213-derived tools for analyzing symbiotic *Vibrio fischeri* reveal patterns of infection and *lux* expression in situ. *Appl. Environ. Microbiol.* **72**, 802–810 (2006).
